# Evolutionary spread of protein L-(iso)aspartyl *O*-methyltransferases guides the discovery of distinct isoaspartate-containing peptides, pimtides

**DOI:** 10.1101/2023.05.11.540355

**Authors:** Hyunbin Lee, Sho Hee Park, Jiyoon Kim, Jaehak Lee, Min Sun Koh, Jung Ho Lee, Seokhee Kim

## Abstract

Ribosomally synthesized and post-translationally modified peptides (RiPPs) are a structurally diverse class of natural products with a distinct biosynthetic logic, the enzymatic modification of genetically encoded precursor peptides. Although their structural and biosynthetic diversity remains largely underexplored, the identification of novel subclasses with unique structural motifs and biosynthetic pathways has been challenging. Here, we report that protein L-(iso)aspartyl *O*-methyltransferases (PIMTs) present in several RiPP subclasses are highly homologous. Importantly, we discovered that the apparent evolutionary transmission of the PIMT gene could serve as a basis to identify a novel RiPP subclass. Biochemical and structural analyses suggest that these homologous PIMTs commonly convert aspartate to isoaspartate via aspartyl-*O*-methyl ester and aspartimide intermediates, and often require cyclic or hairpin-like structures for modification. By conducting homology-based bioinformatic analysis of PIMTs, we identified over 2,800 biosynthetic gene clusters (BGCs) for known RiPP subclasses in which PIMTs install a secondary modification, and over 1,500 BGCs in which PIMTs function as a primary modification enzyme, thereby defining a new RiPP subclass, named pimtides. Our results suggest that the genome mining of proteins with secondary biosynthetic roles could be an effective strategy for discovering novel biosynthetic pathways of RiPPs.

**Insert Table of Contents artwork here:** 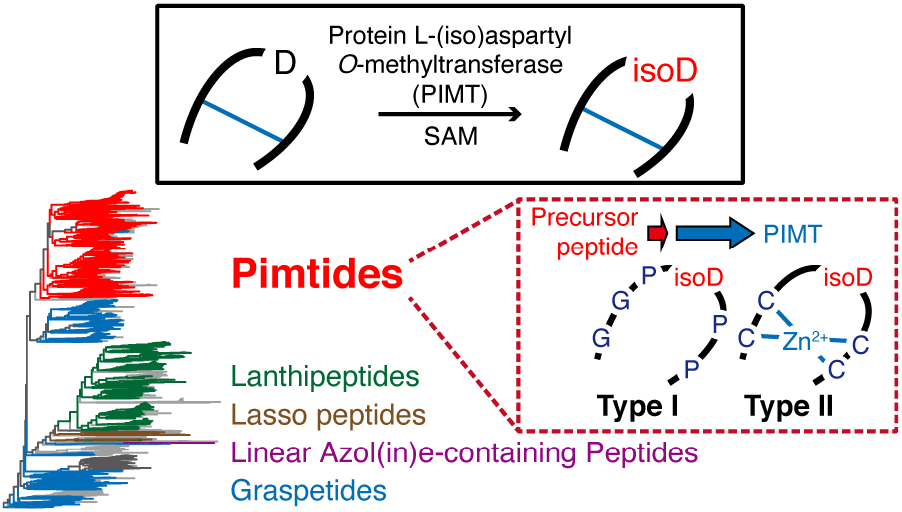

## Introduction

Ribosomally synthesized and post-translationally modified peptides (RiPPs) are a rapidly expanding class of natural products.^1, 2^ RiPPs exhibit a unique biosynthetic pathway that involves genetically encoded precursor peptides and enzymes, with the latter installing diverse post-translational modifications (PTMs) on the precursor peptides. The precursor peptides typically have two modules: the leader and core regions, which are recognized and modified by PTM enzymes, respectively. This biosynthetic simplicity and modularity, and thereby high evolvability, underlies not only the structural and functional diversity of RiPPs but also the high potential for engineering novel functional biomolecules.

The hallmark of RiPP diversity is the distinct PTMs and associated biosynthetic pathways, which individually define the subclasses of RiPPs. The number of RiPP subclasses has increased from 23 in 2013^1^ to 41 in 2021^2^, highlighting the recent expansion of the RiPP diversity, which is likely to continue in the near future. Discovery of new RiPP subclasses has traditionally relied on fortuitous isolation by activity-based screening.^1^ While the explosion of genome sequence data and development of genome mining tools have allowed the identification of a large number of putative biosynthetic gene clusters (BGCs) for RiPPs^3-5^, these methods are generally inefficient at uncovering novel RiPP subclasses; unlike nonribosomal peptide synthetases (NRPSs), RiPP biosynthetic enzymes lack common features across all RiPP subclasses, and therefore, the typical genome mining approach based on homology of biosynthetic enzymes is mostly limited to the expansion of known or closely related RiPP subclasses.

Dissecting phylogenetically distinct orphan BGCs has proven effective for revealing novel PTMs and RiPP-associated pathways, exemplified by recent discoveries of spliceotides^6^, ranthipeptides^7^, streptides^8^, pearlins^9^, and triceptides^10^. However, this approach depends on functional divergence of enzymes known for other class-defining PTMs, and has particularly been successful with an enzyme family with versatile activities, radical *S*-adenosylmethionine (rSAM) enzymes.^11-14^ Furthermore, it is often challenging to find candidate BGCs that significantly diverge from those responsible for known PTMs. To bypass the requirement for homologous biosynthetic enzymes, a marker-independent strategy called decRiPPter has recently emerged.^15^ This method uncovered a new sub-family of lanthipeptides, but it may be biased on the characteristics of known RiPP precursors and requires further exploration to unveil novel PTMs. Overall, new strategies are needed to systematically uncover the unexplored chemical and biosynthetic space associated with RiPPs.

Here, using protein L-(iso)aspartyl *O*-methyltransferase (PIMT) enzymes, we demonstrate that the genome mining of a secondary modification enzyme can promote the identification of a novel RiPP subclass. PIMTs, also known as L-isoaspartyl protein carboxyl methyltransferases (PCMs), were originally discovered as repair enzymes for damaged proteins. They catalyze the conversion of isoaspartate (isoAsp), which is spontaneously formed from an Asp or Asn residue, to Asp (Figure 1A).^16,17^ Recently, PIMT homologs were found in BGCs of several RiPP subclasses, including lanthipeptides, lasso peptides, and graspetides, where they install secondary modifications on conserved Asp residues.^18-22^ This results in the formation of an aspartimide or isoAsp, the former of which is eventually hydrolyzed to the latter (Figure 1).

**Figure 1.**
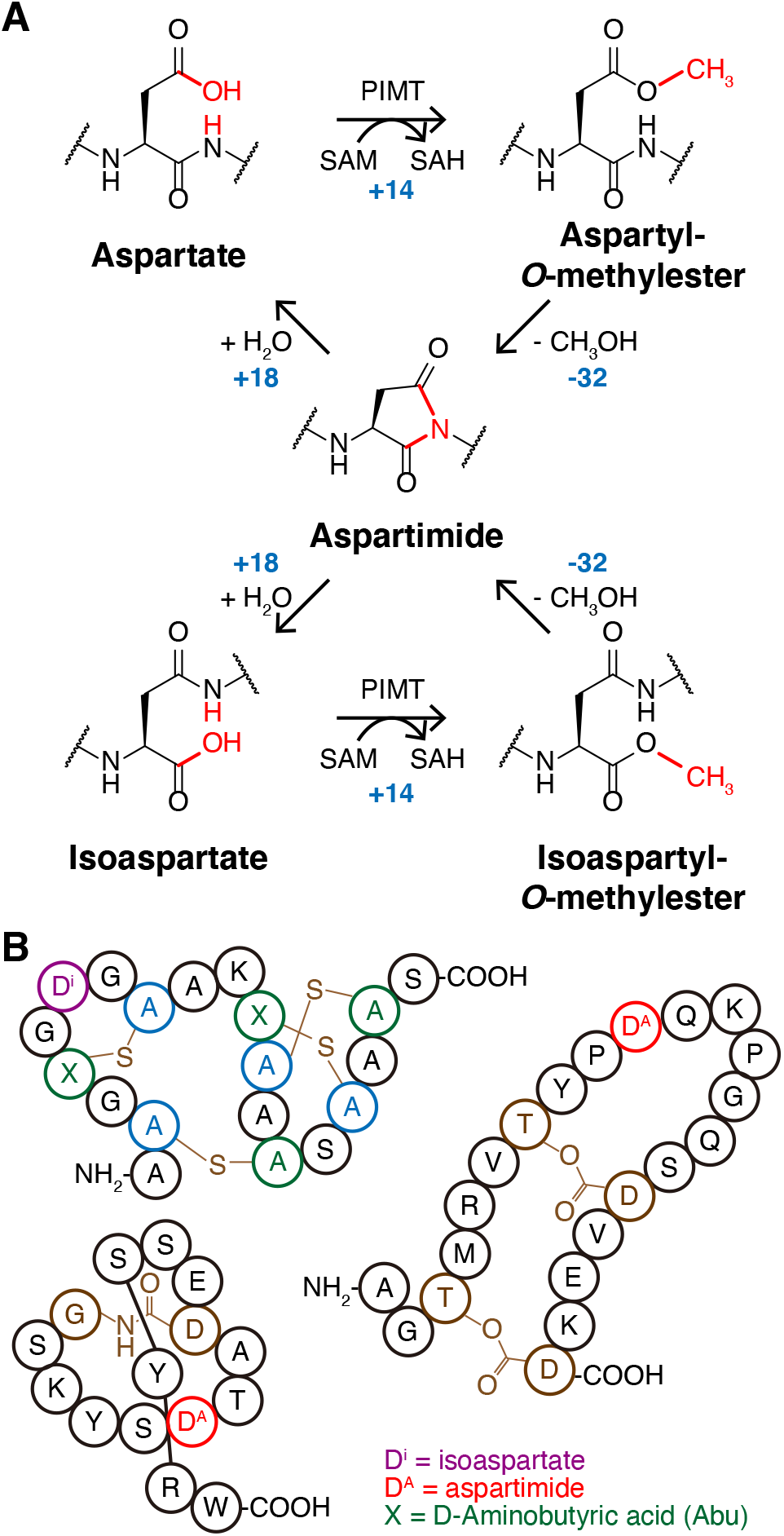
Overview of PIMT-associated peptide maturation. (A) Molecular mechanism of peptide modification by PIMTs. Aspartates (top, left) or isoaspartates (bottom, left) are methylated by PIMT enzymes to yield aspartyl-*O*-methylesters (top, right) or isoasparty-*O*-methylesters (bottom, right). Nitrogen in the backbone amide attacks the carbonyl group to yield aspartimides (middle), and these can be hydrolyzed either to aspartates or isoaspartates. Accompanying molecular weight changes in these conversions are written in blue. (B) Structure of PIMT-modified lanthipeptide (OlvA(BCS)^GluC^, top left), lasso peptide (cellulonodin-2, bottom left), and graspetide (fuscimiditide, right). Isoaspartate and aspartimides are highlighted in purple and red, respectively. D-Amino-butyric acids in the lanthipeptide is denoted as “X” and colored in green. Class-defining modifications are colored in brown.

We have expanded the structural diversity of the graspetide family of RiPPs, also known as omega-ester-containing peptides (OEPs).^23-26^ Graspetides and their biosynthesis have been also studied by bioinformatic^27, 28^, biochemical^29-31^, and structural analysis^32-34^. Using heterologous co-expression, *in vitro* reconstitution of enzyme reactions, and structural analysis of the product peptides with mass spectrometry and NMR, we discovered that SsfM, a new PIMT enzyme associated with graspetide biosynthesis, has the same activity as other PIMT enzymes in RiPP biosynthesis. Genome mining of PIMTs revealed not only homologous enzymes playing a secondary role for known RiPP subclasses, but also those mediating the same Asp-to-isoAsp conversion as the primary modification, leading to a new RiPP subclass. This result highlights the potential of leveraging the evolutionary dissemination of tailoring enzymes across diverse RiPP BGCs to identify a novel biosynthetic pathway of RiPPs.

## Results and Discussion

### *O*-methyltransferase-associated gene cluster produces a new group of graspetides

Our previous genome mining of graspetides focused on identifying a conserved core motif that confidently defines a subgroup of graspetides, leaving room for novel graspetides beyond the twelve groups.^26^ We further inspected homologous ATP-grasp genes and their neighboring genes using Position-Specific Iterative BLAST (PSI-BLAST)^35^ and Rapid ORF Description and Evaluation Online (RODEO)^36^. We found that a large number of ATP-grasp genes (1,288 out of 7,834), mainly from a single clade in a maximum likelihood tree, constitute a distinct BGC that encodes a putative precursor and a PIMT homolog (Figure S1A and 2A). Putative precursor peptides exhibit diverse patterns of the conserved sequence, in which the C-terminus often has an Asp-rich region preceded by a Thr-rich region, suggesting various ω-ester ring connectivities (Figure S1B and Supplementary Dataset 2). Recently, Mitchell and coworkers also reported a comprehensive genome mining of graspetides and designated these putative PIMT-associated RiPPs as group 13 graspetides.^27^

To verify that these BGCs indeed generate graspetides, we chose a BGC from *Streptomyces sp*. F-3 as a model, which encodes a precursor peptide (SsfA), an ATP-grasp enzyme (SsfB), and a PIMT homolog (SsfM; Figure 2A). SsfA and SsfB were co-expressed heterologously in *Escherichia coli*, and the SsfB-modified SsfA, named SsfA(B), was found to be 90 Da lighter than SsfA, indicating the formation of five ester/amide linkages (Figure 2B). To further characterize these linkages, we prepared a GluC-digested SsfA(B) fragment, SsfA(B)_63–97_, and treated it with methoxide, which selectively opens the ω-esters and generates the methyl ester of the ring-forming acidic residue.^26^ We observed up to five-fold methanolysis, suggesting that SsfA(B)_63–97_ has five ester linkages (Figure 2C). The tandem mass (MS/MS) analysis of the five-fold methanolized SsfA(B)_63–97_ suggested that the five ring-forming carboxylates reside within the C-terminal 12 residues, which have only four Asp residues and no Glu (Figure 2D). This result strongly suggests that the C-terminal carboxylate also participates in the ester formation, presenting the first example of the side-to-end linkage in graspetides. Various efforts to determine the ring connectivity using the previously established MS/MS analysis of reaction intermediates or partially hydrolyzed products were unsuccessful.^24-26^ Collectively, these data suggest that the PIMT-containing BGCs produce graspetides. While this manuscript was in preparation, Link and co-workers recently confirmed the graspetide synthesis from homologous BGCs.^20, 21^

**Figure 2.**
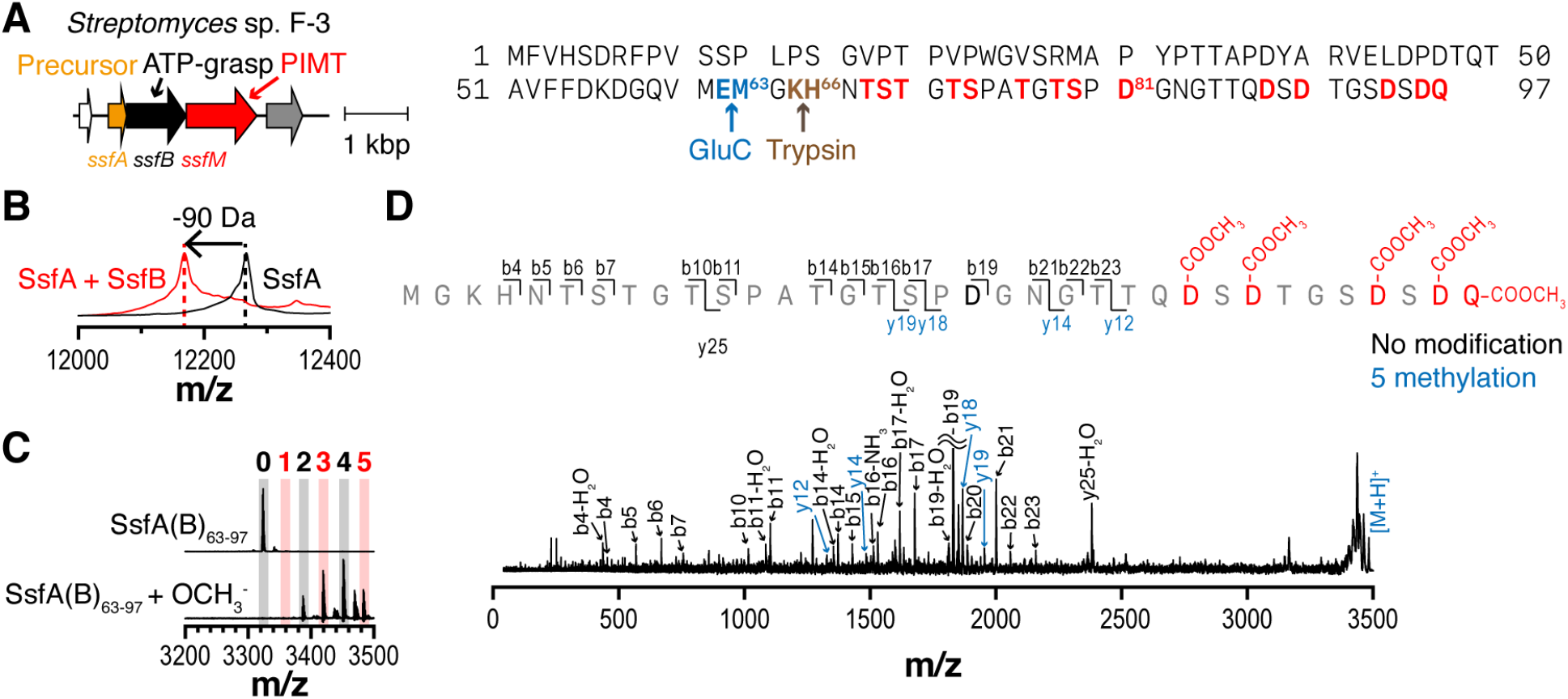
Five ester bonds are generated in the modified peptide. (A) Selected model BGC (left) and precursor peptide sequence (right). Arrows are color-coded based on the predicted domains in the proteins. Cleavage sites by either GluC or trypsin are designated by arrows under the precursor sequence. Residues with hydroxyl or acidic side chains are highlighted in red. (B) MALDI-TOF-MS spectra of SsfA and SsfA(B). (C) MALDI-TOF-MS spectra of GluC-digested SsfA(B) and its methanolyzed product. The numbers of added methanol molecules are specified above each peak. (D) MALDI-TOF-MS/MS spectrum of fully methanolyzed SsfA(B)_63–97_. Observed ions are colored according to the numbers of added methyl groups in each fragment (black, 0; blue, 5). Calculated and observed mass values can be found in **Supplementary Dataset 3**.

### SsfM and other PIMT homologues in RiPP biosynthesis have a common activity

PIMT homologs found in BGCs of lanthipeptides, lasso peptides, and graspetides mediate the conversion of Asp to aspartimide or isoAsp.^18-22^ In a conserved biosynthetic mechanism, they initially methylate the carboxyl side-chain of Asp or iso-Asp to produce (iso)aspartyl-*O*-methyl ester, which is nonenzymatically converted to aspartimide and finally hydrolyzed to isoAsp or Asp (Figure 1A). Biochemical analyses suggest that SsfM also mediates this enzymatic reaction.

First, we co-expressed SsfM with SsfA and SsfB, the modified SsfA, named SsfA(BM), and found that the resulting SsfA(BM) had the same molecular weight (MW) as SsfA(B) (Figure S2A). Unlike other PIMT-modified RiPPs^18-22^, we could not separate the trypsin-digested fragments, SsfA(BM)_63– 97_ and SsfA(B)_63–97_, using HPLC (Figure S2B). Second, we reconstituted the reaction *in vitro* with the purified SsfM, the leader-less SsfA(B)_63–97_, and *S*-adenosylmethionine (SAM). We monitored the reaction at multiple time points by MALDI-TOF-MS and observed an initial gain of 14 Da, followed by a loss of 32 Da and a gain of 18 Da (Figure 3A). These MW changes are consistent with the previously reported reaction pathway of PIMT homologs that includes methylation, aspartimidylation, and hydrolysis (Figure 1A).^18-22^ This result also suggests that the SsfM-mediated reaction does not require the leader region of the precursor peptide, as demonstrated with OlvS, TceM, and LihM.^18, 19, 22^ Third, we isolated the reaction intermediates enriched in the aspartyl-*O*-methyl ester or aspartimide intermediate (+14 Da or -18 Da from the SsfA(B)_63–97_ MW, respectively) and found that they underwent the same MW changes in the absence of SsfM (Figure 3B). This result suggests that the last two steps, aspartimidylation and hydrolysis, do not require SsfM.

**Figure 3.**
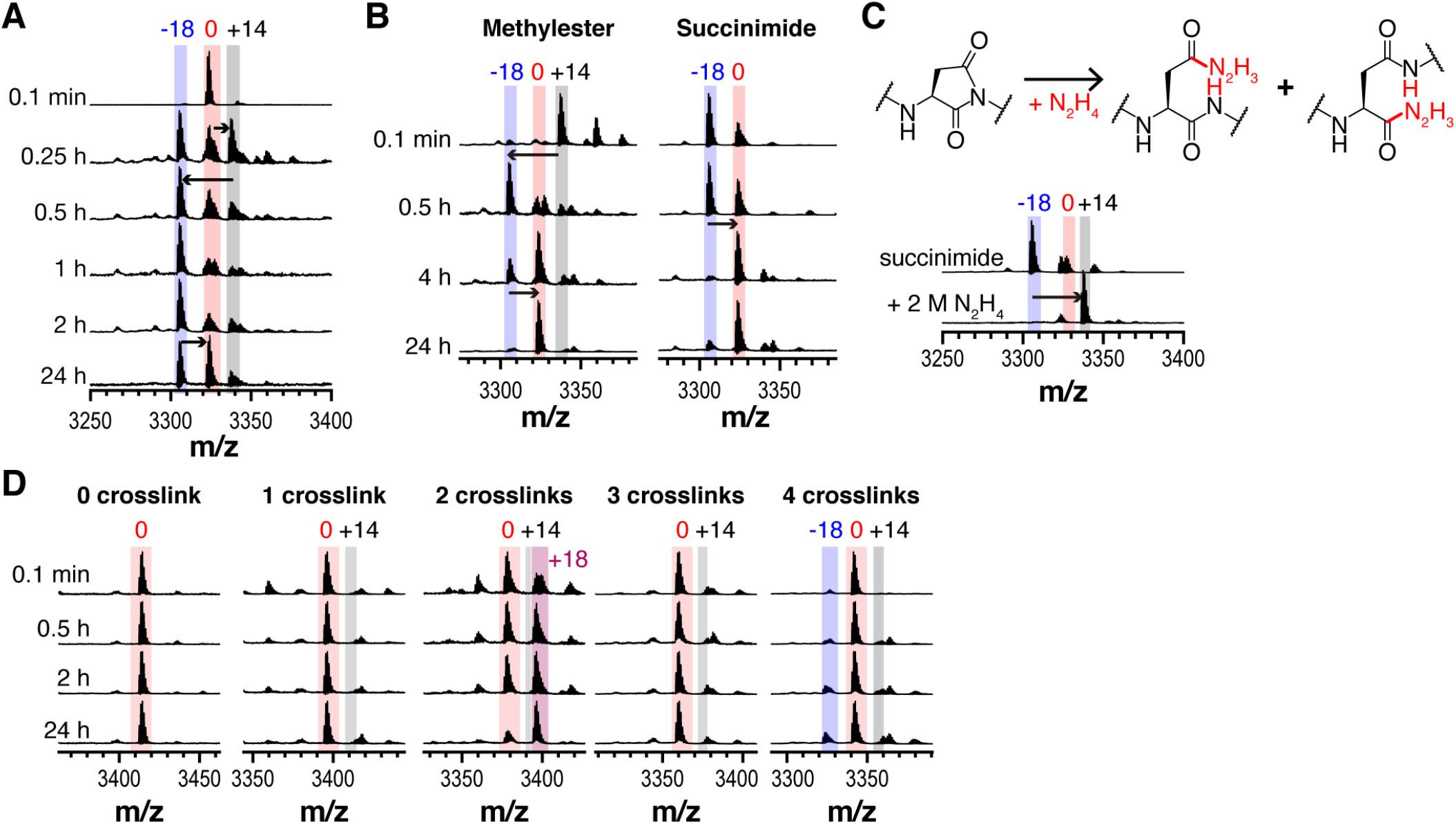
SsfM modifies SsfA(B). (A) MALDI-TOF-MS spectra of the SsfM-mediated modification of SsfA(B)_63–97_ in vitro. Substrates (20 μM) were mixed with SsfM (5 μM) in the presence of SAM (1 mM), DTT (1 mM), and Tris-HCl pH 8.0 (50 mM) at 25 °C. Reactions were quenched at designated time points and monitored by mass analyzer. Relative mass values to SsfA(B)_63–97_ are indicated above peaks. (B) MALDI-TOF-MS spectra of the purified methylester-(left) or succinimide-containing reaction intermediate (right). Purified intermediates were dissolved in 10 mM Tris-HCl pH 8.0 to a final concentration of 10–50 μM at 25 °C. Solutions were monitored by mass analyzer at each time point. (C) MALDI-TOF-MS spectra of succinimide-(top) and hydrazide-containing peptide (bottom). Succinimide-containing peptide was dissolved in 2 M N_2_H_4_ pH 8.0 at a final concentration of 20 μM and incubated at 37 °C for 2 hours. Relative mass value changes to SsfA(B)_63–97_ are given above each peak. (D) MALDI-TOF-MS spectra of SsfM-mediated modification reactions of partially crosslinked SsfA(B)_63–97_ variants. SsfA_63–97_ containing 0-4 ester bonds were incubated with SsfM under the same condition as in (B). Co-purified peptides with one ester crosslinking in the spectra for those with two ester bonds are annotated as +18 Da. Relative mass value changes to the starting materials are given above peaks. Calculated and observed mass values can be found in Supplementary Dataset 3.

Fourth, the nucleophilic addition of hydrazine to the electrophilic aspartimide intermediate resulted in a 32 Da increase, providing further evidence for the presence of the aspartimide intermediate (Figure 3C).^37^ MS/MS analysis of the hydrazine-added and ester-hydrolyzed peptide indicated that Asp81 is the modification site for SsfM (Figure S3). Finally, we explored the substrate scope of SsfM by using partially crosslinked SsfA_63-97_ variants containing 0-4 ester linkages. SsfM did not efficiently modify these partially crosslinked core peptides or the full-length SsfA, indicating that SsfM requires the fully cyclized (five-fold) peptide (Figure 3D and S4). This result is consistent with previous reports^18-22^, suggesting that the PIMT homologs in RiPP biosynthesis generally recognize the cyclized peptide as their substrate. Taken together, these data demonstrate that SsfM shares the same enzymatic activity as other PIMT homologs involved in RiPP biosynthesis.

### SsfM converts Asp81 to isoAsp81

To characterize the SsfM-mediated modification, we enriched SsfA(B)_66–97_ and SsfA(BM)_66–97_ with ^13^C and ^15^N, and analyzed them using multidimensional NMR methods. Initially, we performed 2D ^1^H-^15^N HSQC, 3D HNCACB^38^, 3D HNco-CACB^39^, 3D HNCO^40^, and 3D HNcaCO^41^ experiments to assign the chemical shifts of backbone amide proton (H^N^), nitrogen (N), CO, C^α^, and C^β^ nuclei for all non-proline residues. Uppercase and lower-case letters in the experiment names indicate nuclei with and without frequency labeling, respectively. These analyses identified seven independent peptide fragments that are disconnected at two Pro residues (chain A, B, A’, and B’, Thr68–Ser73; chain C and C’, Ala75–Ser79; chain D and D’, Ala75–Thr78; chain E, F, E’, and F’, Asp81–Gln97 or its smaller part; chain G and G’, Asp81–Asp88) (Figure S5). Multiple NMR resonances originated from a single residue, suggesting the presence of conformational isomers or different chemical structures. All ^1^H-^15^N HSQC resonances in SsfA(B)_66–97_ were found in SsfA(BM)_66–97_, whereas those assigned to smaller regions of chain D’ (Gly77–Thr78) and chain F’ (Asp81–Ser89) were uniquely observed in SsfA(BM)_66–97_ (Figure S5). This result indicates that SsfM mediates additional structural or/and conformational changes on SsfA(B)_66–97_, although the reaction mixture still contained SsfA(B)_66–97_ as a main component. Using a method that can connect the chemical shifts of residues that surround a proline residue, i.e., HNco-cancaNH^42^, we found that chains B, C, and E of SsfA(B)_66–97_ comprise a single long chain (Thr68–Gln97), and chains A and D constitute another chain. Overlapping resonances between SsfA(B)_66–97_ and SsfA(BM)_66–97_ indicate that SsfA(BM)_66–97_ also contains these two long peptides.

Close inspections of spectra clearly indicate the presence of isoAsp at the position of Asp81 in the chain F’ of SsfA(BM)_66–97_. First, the phases of C^α^ and C^β^ cross peaks were inverted both in the strips taken from the ^1^H^N^-^13^C planes at the nitrogen chemical shifts of Gly82 of HNcoCACB and HNCACB spectra (Figure 4). Several studies have previously reported these signals as indicative of the isopeptide linkage.^43-46^ Second, the HNCO signal of Gly82 was not connected to the HNcaCO signal of Asp81, which indicates that C^β^ is between backbone C^α^ and CO, suggesting that isoAsp has been formed (Figure S6). Collectively, the NMR analysis indicates that SsfM converts Asp81 to isoAsp81. We also observed isoAsp at the Asn83 position in chain G of SsfA(B)_66–97_ and in chain G’ of SsfA(BM)_66–97_, suggesting that Asn83 underwent spontaneous aspartimidylation and hydrolysis (Figure S7).^47^

**Figure 4.**
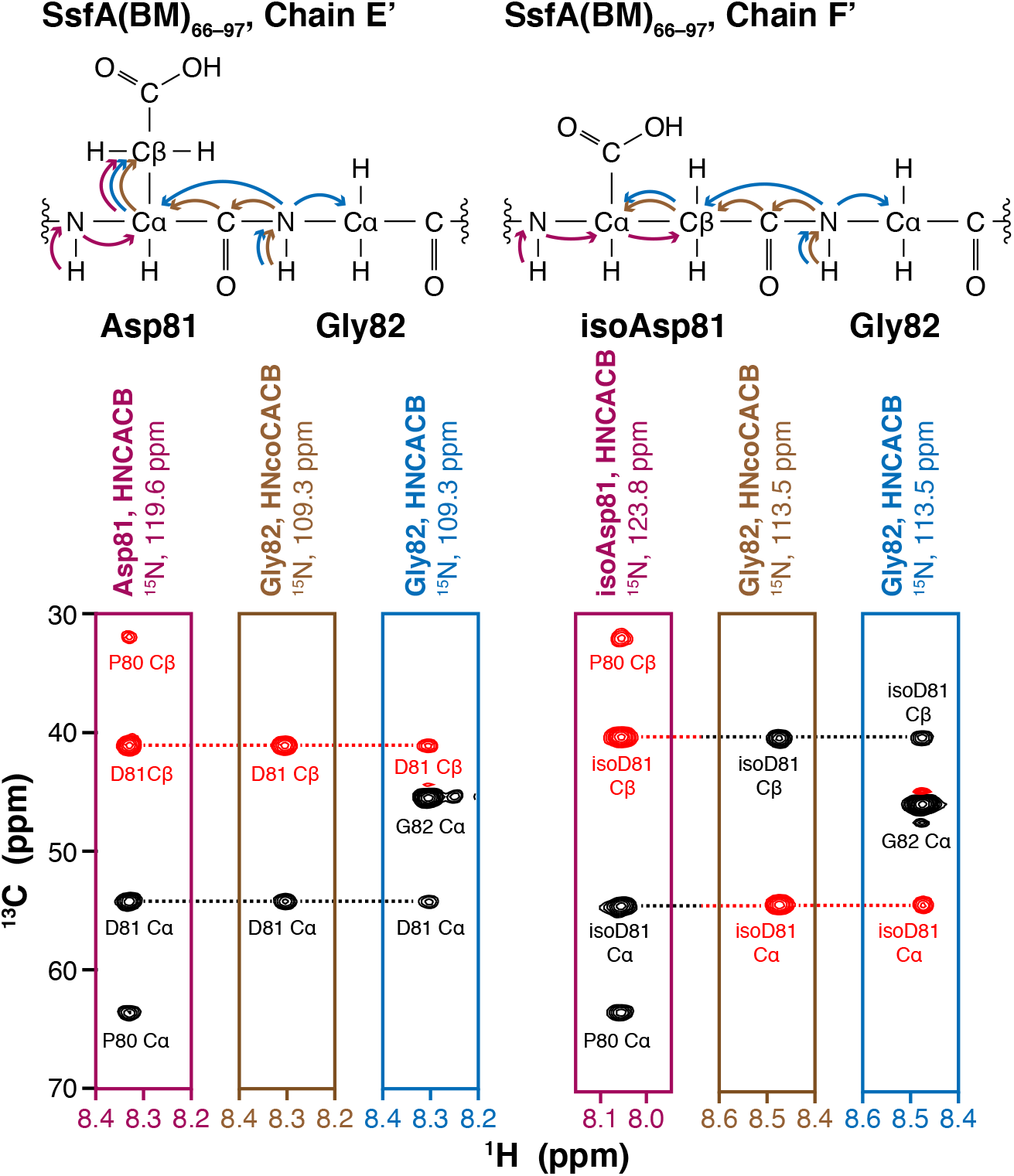
SsfM introduces isoaspartate into SsfA(B). Strip plots of HNCACB and HNcoCACB spectra showing the presence of isoaspartate in SsfA(BM). Magnetization transfer in Asp-Gly and isoAsp-Gly for HNCACB and HNcoCACB experiments are illustrated above. Positive and negative contours are colored in black and red, respectively. Chemical shift values can be found in Supplementary Dataset 3.

### Genome mining of PIMTs reveals a novel RiPP subclass

The common enzymatic activity of PIMTs in the biosynthesis of several RiPP subclasses suggests that these enzymes have close evolutionary relationships. To investigate this further, we used a bioinformatic approach (Figure S8). Initially, we used SsfM as a single query for the PSI-BLAST^35^ to retrieve 73,855 PIMT homologs. To simplify the analysis, we reduced the number of proteins to 23,490 using a cutoff of 70% sequence identity. We generated a maximum likelihood tree and analyzed their domain architecture as well as gene neighbors. We identified putative BGCs for lanthipeptides, lasso peptides, and graspetides, as well as over 1,750 *surE*-*pcm* clusters in which *pcm* encodes a PIMT homolog not associated with RiPP biosynthesis. This PIMT homolog mediates the isoAsp-to-Asp conversion in abnormal proteins in which isoAsp spontaneously arises from Asp and Asn residues, and enhances *E. coli* survival under stress conditions in the late stationary phase.^48^

Notably, we observed that the majority of putative RiPP BGCs were contained in a single clade of 4,003 enzymes, while the *surE*-*pcm* clusters were predominantly located outside of this clade (Figure S8). Additionally, in this clade, 3,200 PIMT homologs (80 %) possessed a C-terminal extension of over 100 amino acids (Figure S8). This C-terminal domain was also identified in PIMTs for lanthipeptides, lasso peptides, and graspetides (OlvS, TceM, and AmdM, respectively).^18, 19, 21^ We used Alphafold^49^ and ColabFold^50^ to obtain predicted 3D structures of SsfM, OlvS, and TceM, and compared them with that of a homologous enzyme from *Thermotoga maritima* (*Tm*PIMT), which is, to our knowledge, the only enzyme in this clade with an experimentally determined 3D structure (PDB 1DL5).^51^ The core structures of their C-terminal domains exhibited a common βαββββα arrangement of secondary structural elements (Figure S9). The pairwise structural alignments on the Dali server^52^ revealed, albeit weak, similarity between these regions with a *Z*-score of 3.4-7.3 (Figure S9). These findings suggest that this domain may have a potential role in RiPP biosynthesis, as recently proposed in a biochemical analysis of a PIMT involved in the maturation of a graspetide, amycolimiditide.^21^

High sequence homology, a conserved C-terminal domain, and the common enzymatic reaction suggest that PIMTs in this clade have evolved from a common ancestor. Furthermore, their frequent association with RiPP biosynthesis implies that this ancestral PIMT has spread to multiple RiPP subclasses. Therefore, we hypothesized that further exploration of this clade could uncover novel RiPP subclasses that utilize PIMTs as either primary or secondary modification enzymes. To test this idea, we compiled the expanded list of PIMTs in this clade without the cutoff of 70% sequence identity and eliminated proteins that are either shorter than 300 amino acids or devoid of genomic information for neighboring genes, resulting in 9,408 enzymes (Figures 5A and S10). Analysis of gene neighbors for known RiPP biosynthetic enzymes or precursor peptides revealed additional BGCs for linear azol(in)e-containing peptides (LAPs; 2 BGCs) as well as lanthipeptides (1,305 BGCs), lasso peptides (67 BGCs), and graspetides (1,432 BGCs; Figure 5A and Supplementary Dataset 2).^2^ Consistent with recent comprehensive genome mining of lanthipeptides^53^, PIMTs in lanthipeptide BGCs are associated with class I lanthipeptides and most precursor peptides adopt the TxDGC core motif (Figure S11A). Precursors for lasso peptides can be classified into two groups based on core motifs; one group contains the DTAD motif in the lasso ring as previously reported^19^, while the other group shares a highly conserved Asp residue in the putative lasso loop (Figures S11B). Precursors encoded in two LAP BGCs have an Asp residue within the C/S/T/G-rich core motif (Figures S11C).^2, 54^ In total, we assigned 2,806 of 9,408 PIMTs into four known subclasses of RiPPs.

**Figure 5.**
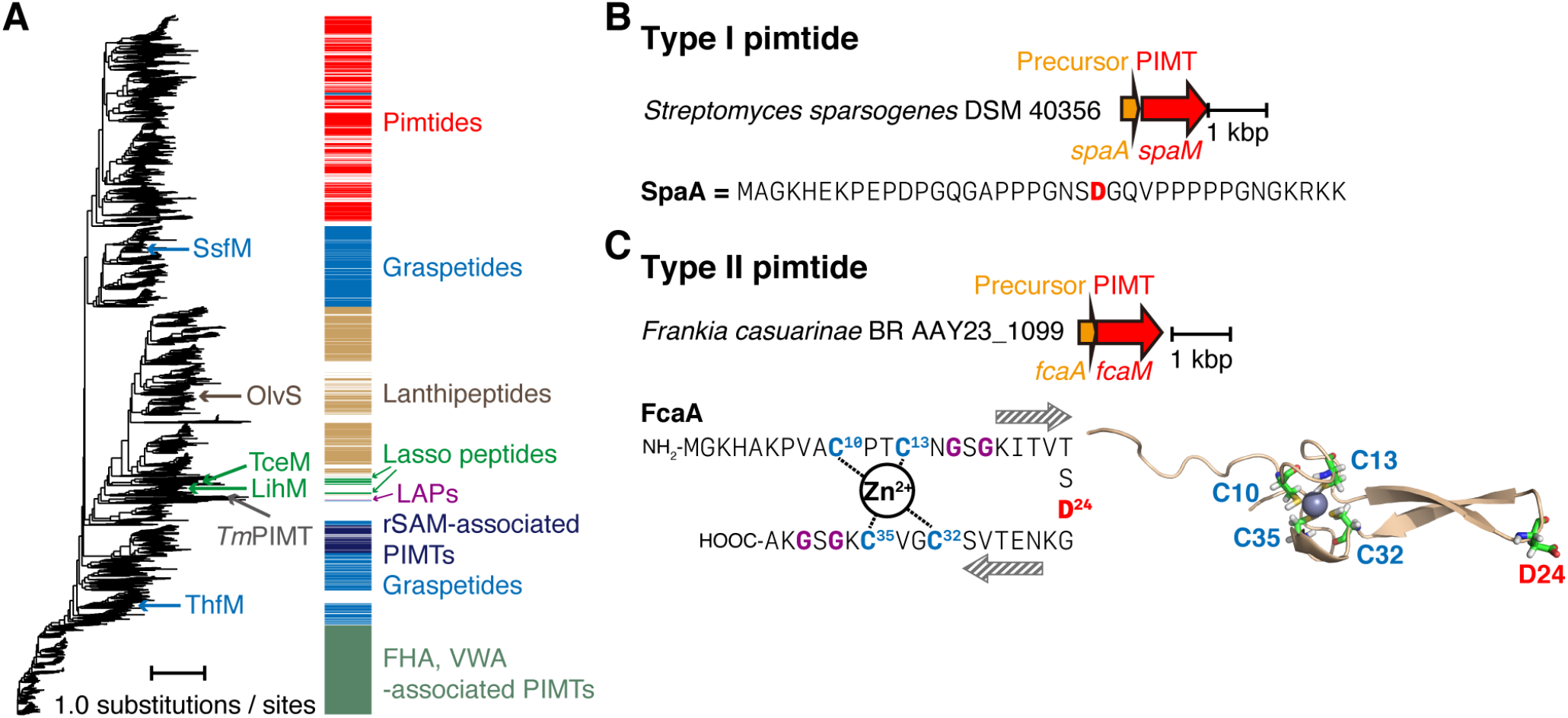
PIMT enzymes are involved in the biosynthesis of various RiPPs. (A) A maximum likelihood tree of 9,408 enzymes is generated, and biochemically characterized PIMTs are annotated. PIMTs encoded in either known RiPP BGCs (graspetides, blue; lanthipeptides, brown; lasso peptides, green; LAPs, purple), newly identified RiPP BGCs (pimtides, red), or conserved gene clusters (FHA-VWA, dark green; rSAM, dark blue) are labeled with color strips. (B-C) Model BGCs for type I pimtides (B; SpaA, precursor; SpaM, PIMT) and type II pimtides (C; FcaA, precursor; FcaM, PIMT). Sequences of precursors (SpaA and FcaA) are shown below each gene cluster. Structure of FcaA was predicted by Alphafold and zinc ion was modelled by MIB. Conserved residues in FcaA are highlighted by colors (cysteine, blue; glycine, purple; aspartate, red).

Nonetheless, most enzymes (70.2 %) did not show any obvious association with known RiPP BGCs. We hypothesized that some of these enzymes could be involved in the biosynthesis of new RiPP subclasses. Indeed, we found a large number of two-gene clusters encoding a putative precursor peptide and a PIMT (1,183 non-redundant putative precursors associated with 1,539 non-redundant PIMTs), but no primary modification enzymes for known RiPPs. Analysis of putative precursor peptides revealed two major types with distinct sequence features. Type I precursors have approximately 45 amino acids and are rich in Gly (17.3 %), Pro (16.8 %), and Asp (10.2 %; Figures 5B and S12A). They present several different conservation patterns of the sequences, but commonly have at least one highly conserved Asp residue nearby conserved prolines or glycines. By contrast, type II precursors contain a highly conserved zinc ribbon motif commonly found in DnaJ (PF00684) with a conserved Asp at the center (CxxCxGxG_D_CxxCxGxG; Figures 5C and 12B).^55^ Four conserved cysteines in the zinc ribbon motif coordinate a zinc ion and the intervening residues form two anti-parallel β-strands (Figure S13). The predicted structure of a type II precursor by Alphafold and a metal ion-binding site prediction server (MIB) also showed the typical zinc ribbon, in which the conserved Asp residue is located in the hairpin (Figure 5C).^49, 56^ The two-gene architecture with a highly homologous PIMT enzyme and the presence of a conserved Asp residue in putative precursors suggest that these PIMTs serve as a primary modification enzyme for the Asp derivatization in the putative precursor, defining a novel subclass of RiPPs. We propose the name “pimtides” for those produced by these distinct BGCs.

We also identified 1,410 PIMTs associated with the conserved gene clusters that typically contain ten genes as well as ABC transporter genes (Figure S14). In particular, the PIMT gene is located next to a gene encoding forkhead-associated (FHA) domain-containing protein. This protein contains a long N-terminal Pro/Gly-rich region with a few Asp residues, similar to the putative precursors for type I pimtides, suggesting that PIMT in this gene cluster may modify the FHA domain-containing protein. We also observed additional 249 distinct gene clusters consisting of two genes encoding a radical *S*-adenosylmethionine (rSAM) enzyme and a PIMT. However, we could not find any neighboring genes encoding putative precursors or substrate proteins for modification (Figure S14B). Although we could not obtain any clues that the remaining 3,653 enzymes are associated with RiPP biosynthesis or protein PTM, we cannot exclude the possibility that these enzymes are also involved in the same type of modification reactions.

### PIMTs in pimtide BGCs mediate the Asp-to-isoAsp conversion

To test whether PIMTs in the pimtide BGCs catalyze the predicted Asp-to-isoAsp conversion in the precursor peptides, we initially selected one BGC for type II pimtide from *Frankia cauarinae* BR AAY23_1099 (Figures 5C and 6A). Heterologous co-expression of the precursor (FcaA) and PIMT (FcaM) in *E. coli* showed that the product, FcaA(M), had the same MW as the unmodified FcaA (Figure S15). We purified FcaA(M) and digested it with trypsin to obtain FcaA(M)_19–26_ (**3**, Figure 6B) containing the conserved Asp residue. We also chemically synthesized the FcaA_19–26_ equivalent (ITVTSDGK, **1**) and its isoAsp variant (ITVTS(isoD)GK, **2**). The comparison of HPLC chromatograms of individual peptides or their combinations revealed that the major component of FcaA(M)_19–26_ is equivalent to the isoAsp variant and clearly different from FcaA_19–26_ (Figure 6B).

**Figure 6.**
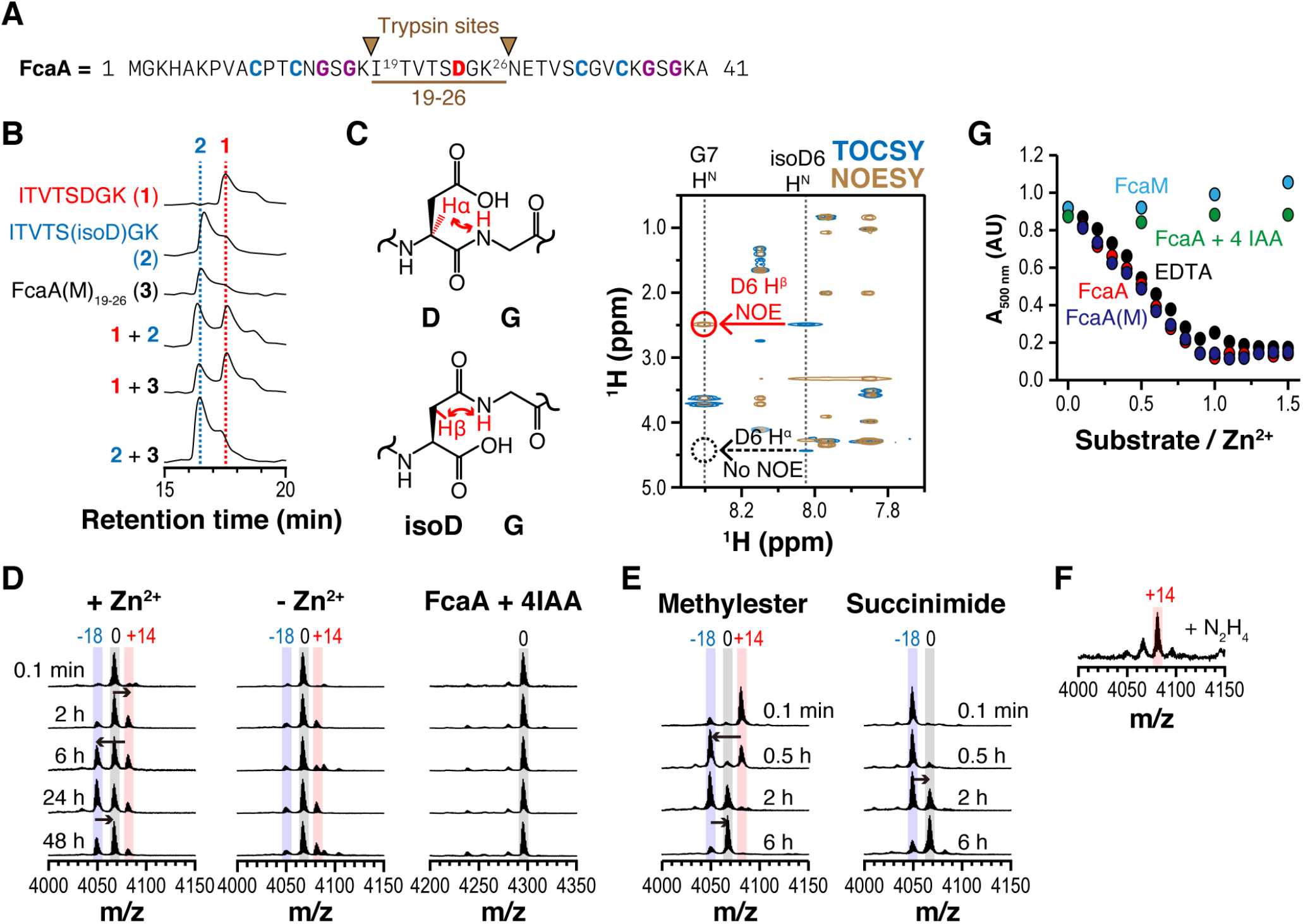
FcaM-mediated isoaspartate installation for a type II pimtide. (A) Trypsin cleavage sites in FcaA. (B) HPLC analysis of FcaA(M)_19-26_ with synthetic peptides **1** (ITVTSDGK) and **2** (ITVTSisoDGK). (C) NMR analysis of FcaA(M)_19-26_. The NOE signal indicates that Asp24 is converted to isoaspartate. (D) MALDI-TOF-MS spectra of FcaM-mediated modification of FcaA or its variant *in vitro*. FcaA or iodoacetamide-labeled FcaA (FcaA+4IAA, 50 μM) was mixed with FcaM (10 μM) in the presence of DTT (1 mM), Tris-HCl pH 8.0 (20 mM), and ZnCl_2_ (0 or 100 μM) at 25 °C. Reactions were monitored at designated time points by mass analyzer. (E) MALDI-TOF-MS spectra showing spontaneous chemical transformation of methylester-(left) or aspartimide-containing intermediates (right). Peptides (20–100 μM) were dissolved in a buffer containing Tris-HCl (20 mM, pH 8.0), DTT (1 mM), and ZnCl_2_ (100 μM). The mixture was incubated at 25 °C for designated time points. (F) MALDI-TOF-MS spectrum of the hydrazide-containing peptide. The reaction condition for Figure 3D was adopted to trap the aspartimide by hydrazine. (G) Competition assay of 4-(2-pyridylazo)resorcinol (PAR) and various substrates toward Zn^2+^. ZnCl_2_ (10 μM) was mixed with PAR (100 μM) in a buffer containing Tris-HCl (20 mM, pH 8.0) and NaCl (100 mM). 0–15 μM substrates were added to the mixture and absorbance at 500 nm was monitored in each mixture. Each data point is colored based on the substrate. Chemical shift values, observed and calculated mass values can be found in Supplementary Dataset 3.

We also obtained the ^1^H, ^1^H-^1^H COSY, ^1^H-^1^H TOCSY, and ^1^H-^1^H NOESY spectra for the three peptides and assigned the chemical shifts of protons. In the NOESY spectrum of FcaA(M)_19-26_, we observed a NOE signal between G25 H^N^ and D24 H^β^, but not between G25 H^N^ and D24 H^α^ (Figures 6C and S16), which is consistent with the previous observation for iso-Asp-containing lanthipeptide OlvA(BCS_A_)^GluC^.^18^ The chemically synthesized ITVTS(isoD)GK (**2**) also showed this correlation, but the unmodified peptide ITVTSDGK (**1**) presented the reverse correlation (Figures S17). These analyses consistently support that FcaM mediates the Asp-to-isoAsp conversion in FcaA.

To characterize the FcaM-mediated reaction in detail, we reconstituted the reaction *in vitro* under various conditions (Figure 6D). In the presence of Zn^2+^, FcaA displayed the same MW changes as those of SsfA(B)_63-97_ and other cyclized intermediates of RiPPs associated with PIMT enzymes: an initial gain of 14 Da, followed by a loss of 32 Da and a gain of 18 Da (Figure 6D). The isolated intermediates also showed spontaneous conversions to the species with the original MW and hydrazine trapping of the putative aspartimide species (−18 Da) resulted in the hydrazine-added species (+14 Da; Figures 6E-F). However, the lack of Zn^2+^ or the iodoacetamide (IAA) labeling of cysteines in FcaA resulted in no MW changes (Figure 6D), indicating that the cysteine-coordinated Zn^2+^ in FcaA is essential for the modification. Additionally, the complex of Zn^2+^ and 4-(2-pyridylazo)resorcinol (PAR)^57^, a metal-sensitive colorimetric reagent, was completely dissociated by adding equal amount of FcaA or FcaA(M), but not by FcaM or IAA-labeled FcaA, suggesting that Zn^2+^ binds to FcaA or FcaA(M) in a 1:1 ratio (Figure 6G). These results suggest that, in line with other PIMT-associated RiPPs, the type II pimtides require the ‘cyclic’ or ‘hairpin-like’ architecture for the Asp-to-isoAsp conversion.

We also tested a model precursor (SpaA) and PIMT (SpaM) for type I pimtides from *Streptomyces sparsogenes* DSM 40356 and found that the SpaM-mediated reaction displayed almost the same features as those of PIMTs in RiPP biosynthesis: no apparent MW change upon heterologous co-expression, the same pattern of MW changes in the *in vitro* reaction, the spontaneous conversions of the reaction intermediates, and the hydrazine trapping of the aspartimide intermediate (Figures 5B and S18). Although we could not directly confirm the Asp-to-isoAsp conversion with SpaM, the same reaction features and the evolutionary relationship of SpaM with other PIMT enzymes suggest that SpaM most likely mediates the same modification reaction. It is unclear whether the type I pimtides also display a cyclic architecture for the modification reaction, but it is possible that the Pro/Gly-rich motifs in type I precursors spontaneously form a cyclic or hairpin-like structure. Collectively, these data suggest that PIMTs in the pimtide BGCs serve as a primary modification enzyme by mediating the same chemical conversion as other PIMT enzymes in RiPP biosynthesis.

## Conclusion

Overall, we report that highly homologous PIMTs have evolutionarily spread across multiple RiPP subclasses and mediate the same modification reaction that converts between Asp and isoAsp via aspartyl-*O*-methyl ester and aspartimide intermediates. More importantly, we show that this evolutionary feature could guide the identification of a novel RiPP subclass, pimtides, in which the PIMT-mediated Asp-to-isoAsp conversion is the primary modification reaction.

Using various biochemical characterizations including heterologous co-expression, *in vitro* reconstitution, time-course experiment, and hydrazine trapping as well as mass spectrometry and NMR analyses, we confirmed the conserved PIMT-mediated Asp-to-isoAsp conversion for a group 13 graspetide and a type II pimtide. We also identified more than 4,300 putative RiPP BGCs encoding a PIMT enzyme by mining bacterial genomes. Currently, PIMT-associated subclasses of RiPPs include pimtides, graspetides, lanthipeptides, lasso peptides, and LAPs. Among them, pimtides are the only subclass in which PIMT functions as a class-defining modification enzyme.

The functional role of the Asp-to-isoAsp conversion is largely unknown. One possibility is that this conversion generates a β-amino acid that has one additional hydrocarbon in the peptide backbone, thus releasing the ring strain in the macrocyclic structures. Indeed, the majority of the characterized PIMTs modify an Asp residue located in the macrocyclic (e.g. graspetides, lanthipeptides, and lasso peptides) or hairpin-like (type II pimtides) region of precursor peptides, requiring fully cyclized peptides as substrates.^18-22^ Another non-exclusive possibility is that this conversion simply changes the structure in the loop or hairpin region, thus diversifying the physical or functional properties of RiPPs.^18^ Alternatively, as previously suggested, the electrophilic aspartimide intermediate might be a functional product^18-22^, although the aspartimide intermediates are generally unstable in a neutral condition (half-life of 1-3 h for modified SsfA or FcaA; Figures 3B and 6E).

The widespread distribution of an accessory protein in multiple RiPP subclasses was also illustrated by the RiPP precursor peptide recognition element (RRE) domain.^58, 59^ It is highly probable that many other proteins with secondary roles in RiPP biosynthesis are also evolutionarily disseminated to unrelated RiPP subclasses. Tailoring enzymes are often shared by distinct classes of natural products^60-63^, and genome mining approach was recently applied to discover unprecedented fungal arginine-containing cyclodipeptides^64^. Given that the genome mining focused on a class-defining modification enzyme typically expands the members of the same RiPP subclass, we believe that genome mining of accessory proteins is a powerful approach to identify novel RiPP subclasses and unprecedented PTMs.

## ASSOCIATED CONTENT

## Supporting Information

## Notes

The authors declare no competing financial interest.

## ABBREVIATIONS

RiPP: Ribosomally synthesized and post-translationally modified peptide;
PIMT: Protein L-(iso)aspartyl *O*-methyltransferase;
MALDI-TOF-MS: Matrix assisted laser desorption/ionization-time of flight-mass spectrometry;
HPLC: High performance liquid chromatography.

