## Supporting Information for "Evolutionary spread of protein L-(iso)aspartyl *O*-methyltransferases guides the discovery of distinct isoaspartate-containing peptides, pimtides"

### Table of Contents


**Materials and Methods** ........ S3

General materials and methods ........ S3
Cloning ........ S3
Heterologous expression and protein purification ........ S3
Size exclusion chromatography ........ S4
Proteolytic digestion ........ S4
Iodoacetamide labeling of FcaA ........ S4
High Performance Liquid Chromatography ........ S4
*In vitro* reconstitution of enzymatic and chemical reactions ........ S4
Solid phase peptide synthesis ........ S4
Assignment of protein backbone chemical shifts of SsfA(BM)$_{66\text{-}97}$ and SsfA(B)$_{66\text{-}97}$ ........ S5
NMR analysis of synthetic peptides and FcaA(M)$_{19\text{-}26}$ ........ S5
Sequential assignment of proline residues in SsfA(B)$_{66\text{-}97}$ ........ S6
Genome mining of graspetides and pimtides ........ S6
Protein structure prediction and analysis ........ S6
Metal binding assay ........ S6

**Figures** ........ S7

Figure S1. Bioinformatic analysis of putative PIMT-modified graspetides. ........ S7
Figure S2. MS and LC analysis of SsfA(B) and SsfA(BM) ........ S8
Figure S3. Mass analysis of ester-hydrolyzed hydrazide-containing SsfA(BM)$_{63–97}$. ........ S9
Figure S4. MALDI-TOF-MS spectra of SsfM-mediated modification reaction of SsfA. ........ S10
Figure S5. Assigned Chains and $^{1}$H-$^{15}$N HSQC spectra of SsfA(B)$_{66–97}$ and SsfA(BM)$_{66–97}$. ........ S11
Figure S6. HNcaCO and HNCO analysis of SsfA(BM)$_{66–97}$ chain E'/F' ........ S13
Figure S7. HNCACB, HNcoCACB, HNCO, and HNcaCO analysis of SsfA(BM)$_{66–97}$ chain G' ........ S14
Figure S8. Bioinformatic analysis workflow of PIMTs ........ S15
Figure S9. Structural analysis of extra C-terminal domains in model PIMTs ........ S16
Figure S10. Genome mining procedure of C-terminal domain-containing PIMTs ........ S17
Figure S11. Features of PIMT-associated RiPPs ........ S18
Figure S12. Sequence logos of precursor peptides for pimtides ........ S19
Figure S13. Structure of zinc ribbon ........ S20
Figure S14. Conserved PIMT-associated gene clusters ........ S21
Figure S15. MALDI-TOF-MS spectra of the purified recombinant FcaA and FcaA(M) ........ S22
Figure S16. NMR spectra of FcaA(M)$_{19–26}$ ........ S23
Figure S17. NMR analysis of synthetic ITVTSDGK and ITVTS(isoD)GK ........ S25
Figure S18. Mass analysis of a precursor-enzyme for type I pimtide ........ S30

**Tables** ........ S32

Table S1. Optimized sequences of precursor peptides, ATP-grasp enzymes, and PIMTs for heterologous expression in *E. coli*. ........ S32
Table S2. Plasmids used in this study. ........ S34
Table S3. Oligonucleotides used in this study. ........ S35

**References** ........ S36

### Materials and Methods

#### General materials and methods

Genes optimized for heterologous expression of recombinant precursor peptides, ATP-grasp enzymes, and PIMTs in *Escherichia coli* (*E. coli*) were synthesized by Gene Universals (USA, Table S1). Oligonucleotides were purchased from Bionics (Korea). All reagents for cloning were purchased from Enzynomics (Korea) unless otherwise specified. *E. coli* ER2566 (DE3) was used for cloning and protein production. Appropriate antibiotics were added to the media and LB-agar plates to a final concentration of 100 μg/mL. Protein concentrations were determined by UV absorbance at 280 nm or 214 nm. Amicon centrifugal filters were purchased from Merck Millipore (USA). Trypsin and chymotrypsin were purchased from Sigma-Aldrich (USA), and endoproteinase GluC was purchased from NEB (USA). For mass analysis, samples were desalted by C18 ZipTip (Merck, USA) if needed, and Bruker UltraFlextreme MALDI-TOF/TOF-MS was used for the analysis (Bruker Daltonics, USA). $^{13}C_6$-d-glucose and $^{15}NH_4Cl$ were purchased from Cambridge Isotopes Laboratorie (USA). Fmoc-protected isoaspartate (Fmoc-Asp-Otbu) was purchased from Alfa Aeser (USA) and the rest of Fmoc-protected amino acids and Wang resin for solid phase synthesis were purchased from GL Biochem (China) unless otherwise noted.

#### Cloning

Plasmids and primers used in this study are listed in Table S2 and S3, respectively. SsfB gene in the dicistronic plasmid pHB665 was deleted by inverse-PCR mutagenesis[1] to obtain a plasmid encoding recombinant SsfA. *E.coli* maltose-binding protein (MBP) and TEV protease cleavage site ([TEV]) were fused to the N-terminus of FcaA and SpaA by overlap-extension PCR[2]. In brief, F_22b and R_TEV were used as primers to generate the $His_6$-MBP-[TEV] fragment, and R_22b and F_XxxA (XxxA = FcaA or SpaA) were used to yield XxxA fragment at the first PCR step. Two fragments were used as template, and R_TEV and F_XxxA were used as primers of the second PCR step.

PCR products were purified by LaboPass PCR Purification Kit (Cosmogenetech, Korea) following the instructor's guide. Purified products were phosphorylated and ligated using T4 Polynucleotide kinase and T4 DNA ligase). Ligated DNA products were transformed into *E. coli*. Cells were grown on an LB-agar plate containing corresponding antibiotics at 37 °C for 12–16 hours. Each colony was inoculated in 2 mL of LB with antibiotics at 37 °C for 12–16 hours, and plasmids were purified using LaboPass Plasmid Miniprep Kit (Cosmogenetech, Korea). Sequences were verified by Sanger sequencing method (Bionics, Korea).

#### Heterologous expression and protein purification

Plasmids encoding desired proteins were transformed into *E. coli*. Cells were grown on an LB-agar plate with appropriate antibiotics at 37 °C for 12-16 hours. A single colony was inoculated in 10–80 mL of LB containing antibiotics and grown at 37 °C for 12–16 hours. The culture was 100-fold diluted in 1–8 L of fresh LB with antibiotics and further inoculated at 37 °C. When $OD_{600}$ reached between 0.4 and 0.6, protein production was induced by adding β-D-1-thiogalactopyranoside (IPTG; LPS solutions, Korea) to a final concentration of 0.1 mM. Cells were incubated at 25 °C for 16–24 hours and then they were harvested by centrifugation at 5,200 xg for 15 minutes. Cell pellets were stored at - 80 °C for less than four days before use.

To obtain the $^{13}C$- and $^{15}N$-labeled SsfA(B) or SsfA(BC), 1 L of M9 media with the following components were prepared for protein production: $^{15}NH_4Cl$ (1.5 g; Cambridge Isotopes Laboratories), $^{13}C$ glucose (2.0 g; Cambridge Isotopes Laboratories), $Na_2HPO_4$ (6.0 g; LPS solutions), $KH_2PO_4$ (3.0 g; LPS solutions), NaCl (0.5 g; LPS solutions), ISOGRO- $^{13}C$, $^{15}N$ powder (0.5 g; Sigma-Aldrich), 100X MEM vitamin solution (5 mL; ThermoFisher, USA), $MgCl_2$ (1 mM at final; LPS solutions), $CaCl_2$ (0.1 mM at final; LPS solutions), and 1,000X trace metal solution (24.7 mM $FeCl_3$, 0.76 mM $CuCl_2$, 0.05 mM $MnCl_2$, 0.77 mM $CoCl_2$, $H_3BO_3$ 1.6 mM, and $ZnCl_2$ 6.16 mM; 1 mL). Single colony harboring pHB665 or both pHB665 and pHB666 were inoculated in 50 mL of LB containing ampicillin or both ampicillin and streptomycin at 37 °C for 16 hours. When $OD_{600}$ was reached around 3.0–4.0, cells were harvested and transferred into 1L of the prepared M9 media to have an initial $OD_{600}$ around 0.2. Cells were grown at 37 °C until $OD_{600}$ reached 0.5–0.6. Protein production was induced by adding IPTG to a final concentration of 0.1 mM. Cells were further incubated at 25 °C for 24 hours and harvested by centrifugation.

Cell pellets were resuspended in 20–160 mL of wash buffer (50 mM Tris-HCl pH 8.0, 20 mM imidazole, 300 mM NaCl) and lysed by sonication. Insoluble fractions were eliminated by centrifugation at 18,000 xg for 30 minutes at 4 °C. Supernatants were applied to 1–5 mL of Ni resin pre-equilibrated with wash buffer (Ni Sepharose 6 Fast Flow beads, GE Healthcare, USA). Resins were washed twice with a volume of wash buffer equal to 5 column volumes. Proteins were eluted by adding 3 column volume of elution buffer (50 mM Tris-HCl pH 8.0, 500 mM imidazole, 100 mM NaCl). Fractions were concentrated and the buffer was exchanged to buffer A (10 mM Tris-HCl pH 8.0, 100 mM NaCl) using Amicon centrifugal filter. Enzymes were further purified by size exclusion chromatography, and precursor peptides were digested by appropriate proteases and purified by HPLC.

**Size exclusion chromatography**

Size exclusion chromatography was performed using ÄKTA pure 25M1 (GE Healthcare, USA) on a Superdex200 10/300 GL column (Cytiva, USA) equilibrated in buffer A. Fractions were monitored by UV absorbance at 280 nm and SDS-PAGE. Desired fractions were collected and concentrated using Amicon centrifugal filter, and stored at –80 °C.

**Proteolytic digestion**

To eliminate SsfB (and SsfC) from SsfA(B) (or SsfA(BC)), trifluoroacetic acid was added to purified SsfA(B) + SsfB (or SsfA(BC) + SsfB + SsfC) to a final concentration of 1 % (v/v), and 1 M Tris-HCl pH 8.0 was added to the mixture to neutralize pH up to 7.0–8.0. Precipitated enzymes were eliminated by centrifugation at 20,000 xg for 10 minutes at 4 °C. Trypsin or GluC was added to the supernatant containing SsfA derivatives (1/100 of SsfA derivatives by weight), and incubated at 37 °C for 16 hours. Cleavage reaction was monitored by MALDI-TOF-MS, and desired fragments were further purified by HPLC.

MBP-fused precursor peptide derivatives (1 mM; FcaA, FcaA(M), SpaA, or SpaA(M)) were digested by adding 10 µM of TEV protease, 1 mM dithiothreitol (DTT), and 20 mM Tris-HCl pH 8.0. The mixtures were incubated at 25 °C for 16 hours. MBP was eliminated by an Amicon centrifugal filter, and the peptides were further purified by HPLC to fully eliminate bound zinc ions from FcaA variants.

To yield FcaA(M)$_{19\text{-}26}$, trypsin was added to the HPLC-purified FcaA(M) (1/10 by weight) and reaction mixture was incubated at 37 °C for 24–48 hours. The reaction was monitored by MALDI-TOF-MS and purified by HPLC.

**Iodoacetamide labeling of FcaA**

The following mixture was prepared to alkylate cysteines in FcaA: 100 µM FcaA, 1 mM iodoacetamide, 1 mM TCEP, 20 mM Bicine pH 8.5. Reaction mixture was incubated at 37 °C for 2 hours. Reactions were monitored by MALDI-TOF-MS and purified by HPLC.

**High Performance Liquid Chromatography**

HPLC was performed using Agilent 1260 Infinity (Agilent, USA). A ZORBAX SB-C18 analytical column (4.6 x 250 mm, particle size 5 µm, Agilent, USA) or a ZORBAX SB-C18 Semi-preparative column (9.4 x 250 mm, particle size 5 µm, Agilent, USA) was used for the LC analysis or purification, respectively. Two mobile phases were used in HPLC: Solvent A (0.05 % Trifluoroacetic acid in $H_2O$, v/v) and solvent B (0.05 % Trifluoroacetic acid in $CH_3CN$, v/v). Peptides were separated by linearly increasing the concentration of $CH_3CN$. Fractions were monitored by UV absorbance at 214 nm or 280 nm, and MALDI-TOF-MS. Desired fractions were collected and freeze-dried. Dried peptides were dissolved in milli-Q $H_2O$ immediately before use.

***In vitro* reconstitution of enzymatic and chemical reactions**

SpaM- or SsfM-mediated modification reaction of SpaA or SsfA variants were reconstituted as follows: 20 µM of substrate, 5 µM of enzyme, 1 mM of S-adenosylmethionine (SAM), 1 mM of DTT, and 50 mM of Tris-HCl pH 8.0. FcaM-mediated reactions of FcaA variants were reconstituted as follows: 100 µM of substrate were mixed with FcaM (20 µM) in presence of DTT (1 mM), Tris-HCl pH 8.0 (20 mM), and $ZnCl_2$ (100 µM). All reaction mixtures were incubated at 25 °C and monitored by MALDI-TOF-MS at designated time points. Reaction mixtures were subjected to HPLC to purify the observed intermediates.

To observe the spontaneous transformation of aspartyl-*O*-methylester to aspartimide and aspartimide to aspartate/isoaspartate, peptides containing aspartyl methylester or aspartimide were dissolved in either 10 mM Tris pH 8.0 (SsfA or SpaA derivatives) or 20 mM Tris pH 8.0, 1 mM DTT, and 100 µM $ZnCl_2$ (FcaA derivatives) to the final concentrations of 10–100 µM. All reaction mixtures were incubated at 25 °C and monitored by MALDI-TOF-MS at designated time points.

For the generation of aspartyl hydrazide, aspartimide-containing peptides were dissolved in 2 M hydrazine pH 8.0 to the final concentrations of 20–100 µM. Solutions were incubated at 37 °C for 2 hours. Reactions were monitored by MALDI-TOF-MS and MALDI-TOF/TOF-MS, if needed. For the SsfA derivative, NaOH was added to a final concentration of 0.1 M to hydrolyze ester bonds.

**Solid phase peptide synthesis**

Peptides (ITVTSDGK and ITVTS(isoD)GK) were synthesized as previously described with modifications.[3] First amino acid was coupled as follows: Wang resin was mixed with Fmoc-protected lysine (5 equiv.) and 4-Dimethylaminopyridine (DMAP, 0.1 equiv.) in *N*,*N*-dimethylformamide (DMF). *N*,*N*'-Diisopropylcarbodiimide (DIC, 4 equiv.) was added to the mixture after 20 minutes. The mixture was further incubated at room temperature for 40 hours.

Remaining amino acids were coupled as follows: 20% piperidine in DMF (v/v) was added to the resins to remove Fmoc groups. Mixtures were incubated at room temperature for 1 hour. Fmoc-protected amino acid (5 equiv.) was dissolved in DMF containing HATU (5 equiv.) and *N*,*N*'-Diisopropylethylamine (DIPEA, 10 equiv.) and mixed with the resins to attach the amino acid. The mixture was incubated at room temperature for 1 hour. Resins were washed with DMF and dichloromethane three times between each step, and each reaction was incubated under rigorous shaking. The coupling reaction was finished by removing Fmoc group.

Cleavage cocktail (TFA:$H_2O$:Triisopropylsilane = 950:25:25, v/v) was added to the resins to detach peptides and remove the remaining protecting groups. The mixture was incubated for 2 hours. Peptides were concentrated by evaporating TFA through air-drying. Ice-cooled ether/hexane (1:1, v/v) was added to the solution to precipitate peptides. Pellets were dried at 50 $^{o}$C, dissolved in DMSO, and purified by HPLC.

**Assignment of protein backbone chemical shifts of SsfA(BM)$_{66-97}$ and SsfA(B)$_{66-97}$**

All NMR experiments were acquired using a Bruker AVANCE NEO 600 MHz NMR spectrometer equipped with a 5 mm triple-resonance z-gradient cryogenic probehead. 0.3 mM $^{13}$C, $^{15}$N-labeled SsfA(BM)$_{66-97}$ and 0.8 mM $^{13}$C, $^{15}$N-labeled SsfA(B)$_{66-97}$ were separately prepared in the 20 mM sodium phosphate buffer (pH 6.0) containing 20 mM sodium chloride and 5% $D_2O$ and placed in a 5 mm $D_2O$-matched Shigemi tube.

Backbone assignments of SsfA(BM)$_{66-97}$ at 277.1 K were carried out by performing 3D HNCACB[4], HNcoCACB[5], 3D HNCO[6], and 3D HNcaCO[7] experiments using the BEST scheme[8]. Small case letters indicate nuclei whose chemical shifts were not recorded. 3D HNCACB data was acquired using a data matrix of 120 ($t_1$, $^{13}$C) × 290 ($t_2$, $^{15}$N) × 1024 ($t_3$, $H^N$) complex points and sweep widths of 10000 ($^{13}$C), 1176 ($^{15}$N), and 6250 ($H^N$) Hz. The total experiment time was 29 h. 3D HNcoCACB data was acquired using a data matrix of 486 ($t_1$, $^{13}$C) × 230 ($t_2$, $^{15}$N) × 1024 ($t_3$, $H^N$) complex points and the same sweep widths as in HNCACB. The total experiment time was 30 h. 3D HNCO data was acquired using a data matrix of 136 ($t_1$, $^{13}$C) × 300 ($t_2$, $^{15}$N) × 1024 ($t_3$, $H^N$) complex points and sweep widths of 1057 ($^{13}$C), 1176 ($^{15}$N), and 6250 ($H^N$) Hz. The total experiment time was 36 h. 3D HNcaCO data was acquired using a data matrix of 148 ($t_1$, $^{13}$C) × 300 ($t_2$, $^{15}$N) × 1024 ($t_3$, $H^N$) complex points and sweep widths of 1359 ($^{13}$C), 1065 ($^{15}$N), and 6250 ($H^N$) Hz. The total experiment time was 39 h. Because the chains in SsfA(B)$_{66-97}$ present in SsfA(BM)$_{66-97}$, the above-mentioned 3D experiments can be used to assign SsfA(B)$_{66-97}$. However, to improve the accuracy of SsfA(B)$_{66-97}$ assignment, an additional 3D HNCACB experiment was performed using the SsfA(B)$_{66-97}$ sample. 3D HNCACB data was acquired using a data matrix of 1024 ($t_1$, $H^N$) × 230 ($t_2$, $^{15}$N) × 486 ($t_3$, $^{13}$C) complex points and sweep widths of 10000 ($^{13}$C), 1176 ($^{15}$N), and 6250 ($H^N$) Hz. The total experiment time was 96 h. A recycle delay of 0.5 s and four scans per increment were used in all 3D assignment experiments. All NMR data were processed by NMRPipe[ref6]. 90°-shifted sine-bell window functions were applied to all dimensions prior to zero-filling and Fourier transformation. All NMR spectra were analyzed by NMRFAM-SPARKY[9].

**NMR analysis of synthetic peptides and FcaA(M)$_{19-26}$**

$^{1}$H, $^{1}$H-$^{1}$H COSY, $^{1}$H-$^{1}$H TOCSY, and $^{1}$H-$^{1}$H NOESY NMR spectra of synthetic peptides and FcaA(M)$_{19-26}$ were acquired at 293.1 K. All samples were dissolved in 600 µL of $d_6$-DMSO at 4.0 mg/mL concentration. All data were processed and analyzed by MestReNOVA[10].

NMR experimental parameters used for synthetic peptides and FcaA(M)$_{19-26}$

| Experiment | Sweep width | Data matrix | Number of scans (per increment) | Mixing time | Total acquisition time |
|---|---|---|---|---|---|
| $^{1}$H | 17857 Hz | 16384 | 256 | - | 13 m |
| $^{1}$H-$^{1}$H COSY | 5882 Hz (in both $^{1}$H dimensions) | 2048 ($t_1$, $^{1}$H) × 256 ($t_2$, $^{1}$H) | 4 | - | 39 m |
| $^{1}$H-$^{1}$H TOCSY | 5882 Hz (in both $^{1}$H dimensions) | 2048 ($t_1$, $^{1}$H) × 1024 ($t_2$, $^{1}$H) | 8 | 80 ms | 5 h 21 m |
| $^{1}$H-$^{1}$H NOESY | 5882 Hz (in both $^{1}$H dimensions) | 2048 ($t_1$, $^{1}$H) × 512 ($t_2$, $^{1}$H) | 8 | 400 ms | 2 h 24 m |

$t_1$ and $t_2$ indicates direct dimension and indirect dimension, respectively.

**Sequential assignment of proline residues in SsfA(B)$_{66\text{-}97}$**

The 4D HNcocancaNH (x-P-x) experiment by Wong et al.[11] correlates two backbone amide groups, i.e., $N_{i+1}H_{i+1}$, and $N_{i-1}H_{i-1}$, that are flanking a proline residue at position i. The 4D experiment was run in a 3D mode without frequency labeling in the $H_{i+1}$ dimension. 0.26 mM $^{13}$C, $^{15}$N-labeled SsfA(B)$_{66\text{-}97}$ was prepared in the 20 mM sodium phosphate buffer (pH 6.0) containing 20 mM sodium chloride and 5% $D_2O$ and placed in a 5 mm $D_2O$-matched Shigemi tube. The 3D hNcocancaNH data was acquired with a recycle delay of 1.5 s, forty scans per increment, a data matrix of 64 ($t_1$, $^{15}$N) × 64 ($t_2$, $^{15}$N) × 2048 ($t_3$, $H^N$) complex points, and sweep widths of 1176 ($^{15}$N), 1176 ($^{15}$N), and 2048 ($H^N$) Hz. The total experiment time was 84 h. The NMR data were processed as described in the protein backbone assignment section.

**Genome mining of graspetides and pimtides**

To identify graspetide biosynthetic gene clusters (BGCs) encoding PIMT, Position-Specific Iterative BLAST (PSI-BLAST)[12] was performed in January 2021 on a non-redundant protein database in National Center for Biotechnology Information (NCBI, http://www.ncbi.nlm.nih.gov) using PsnB as a query and 1e-30 as a cut-off value for four iterations, yielding 7,834 ATP-grasp enzymes. These proteins were aligned by MAFFT 7.453 with the G-large-ins-1 algorithm[13] and the resulting alignment was used to generate a maximum likelihood tree using FastTree 2.1.11[14]. Also, proteins were analyzed using Rapid ORF Description and Evaluation Online (RODEO)[15] to retrieve the information of neighbor genes. ATP-grasp enzymes which are encoded between the genes for a PIMT homolog and a precursor-like peptide were designated as putative PIMT-associated graspetides. These proteins and previously identified enzymes for twelve graspetide groups[16] were annotated in the tree using ETE3 3.1.2[17]. Precursor peptides were analyzed by EFI-EST tool (http://efi.igb.illinois.edu/efi-est) with a cut-off value of 1e-15 to group them by the homology.[18] Sequences of three major groups were aligned by MAFFT 7.453 with E-ins-i algorithm[19] and sequence logos for three major groups were generated based on the alignments using WebLogo3[20].

Bioinformatic analysis for PIMTs are illustrated in Figure S8 and S10. For the retrieval of PIMTs, PSI-BLAST was performed in December 2021 on a non-redundant protein database in NCBI using SsfM as a query and 1e-25 as cut-off value for four iterations, as described in Figure S9. Total 73,855 proteins were obtained, and dereplicated by the sequence identity of 70 %, yielding 23,490 proteins. These proteins were aligned by MAFFT 7.453 with the G-large-ins-1 algorithm[13] and the resulting alignment was used to generate a maximum likelihood tree using FastTree 2.1.11[14]. To predict the biological role of the proteins, proteins were analyzed by RODEO[15] and neighbor genes were analyzed whether they are likely to be involved in the biosynthesis of lasso peptides/lanthipeptides or are likely to be form a *surE-pcm* gene cluster for long-term survival of cells. The tree was visualized by ETE3 3.1.2[17] with their domain architectures and their predicted roles are annotated in parallel.

Finally, 4,003 PIMTs and proteins that were dereplicated by these enzymes were further analyzed as illustrated in Figure S11. First, neighbor genes and their sequences were retrieved using RODEO.[15] PIMTs associated in known RiPP BGCs were classified. Next, short ORFs (≤ 100 amino acids) encoded near PIMT (± 500 bps) and containing aspartate(s) were classified by their homology using a cut-off value of 1e-06 in EFI-EST.[18] Clusters with more than 5 peptides with conserved aspartates were assigned to the putative pimtide BGCs. Finally, proteins encoded in conserved gene clusters containing Forkhead-associated (FHA) domain-containing proteins and von Willebrand factor type A (vWA) domain-containing proteins are assigned to FHA-VWA gene clusters. Proteins were aligned by MAFFT 7.429 with G-large-ins-1 algorithm[13], a maximum-likelihood tree was built with the alignment with FastTree 2.1.11[14], and the tree was visualized by ETE3 3.1.2[17]. Precursor peptides were aligned by MAFFT E-ins-i option[19] and sequence logos were generated using WebLogo3[20].

**Protein structure prediction and analysis**

Structures of SsfM, OlvS, TceM, and FcaA were predicted in Alphafold.[21] To locate the $Zn^{2+}$ ion in the model structure of FcaA, alphafold structure was queried in MIB (http://combio.life.nctu.edu.tw/MIB2/).[22] Structural alignment of SsfM, OlvS, TceM with the crystal structure of PIMT from *Thermotoga maritima* (PDB 1DL5) were generated in DALI server.[23]

**Metal binding assay**

Stoichiometry between binding of zinc ion and peptides/proteins were determined using 4-(2-pyridylazo)resorcinol (PAR)[24] as follows: $Zn^{2+}$-2 PAR complexes were first prepared by mixing 20 µM of $ZnCl_2$, 200 µM of PAR in 20 mM Tris-HCl pH 8.0 and 100 mM NaCl to a final volume of 50 µL. To dissociate the complex, the mixture was 2-fold diluted by adding 50 µL of FcaA, FcaA(M), IAA-labeled FcaA, FcaM, or EDTA. Final concentrations of the added peptides/proteins/EDTA were 1–15 µM. Changes in absorbance at 500 nm was observed using Infinite M200 pro (Tecan, Switzerland).

#### Figures

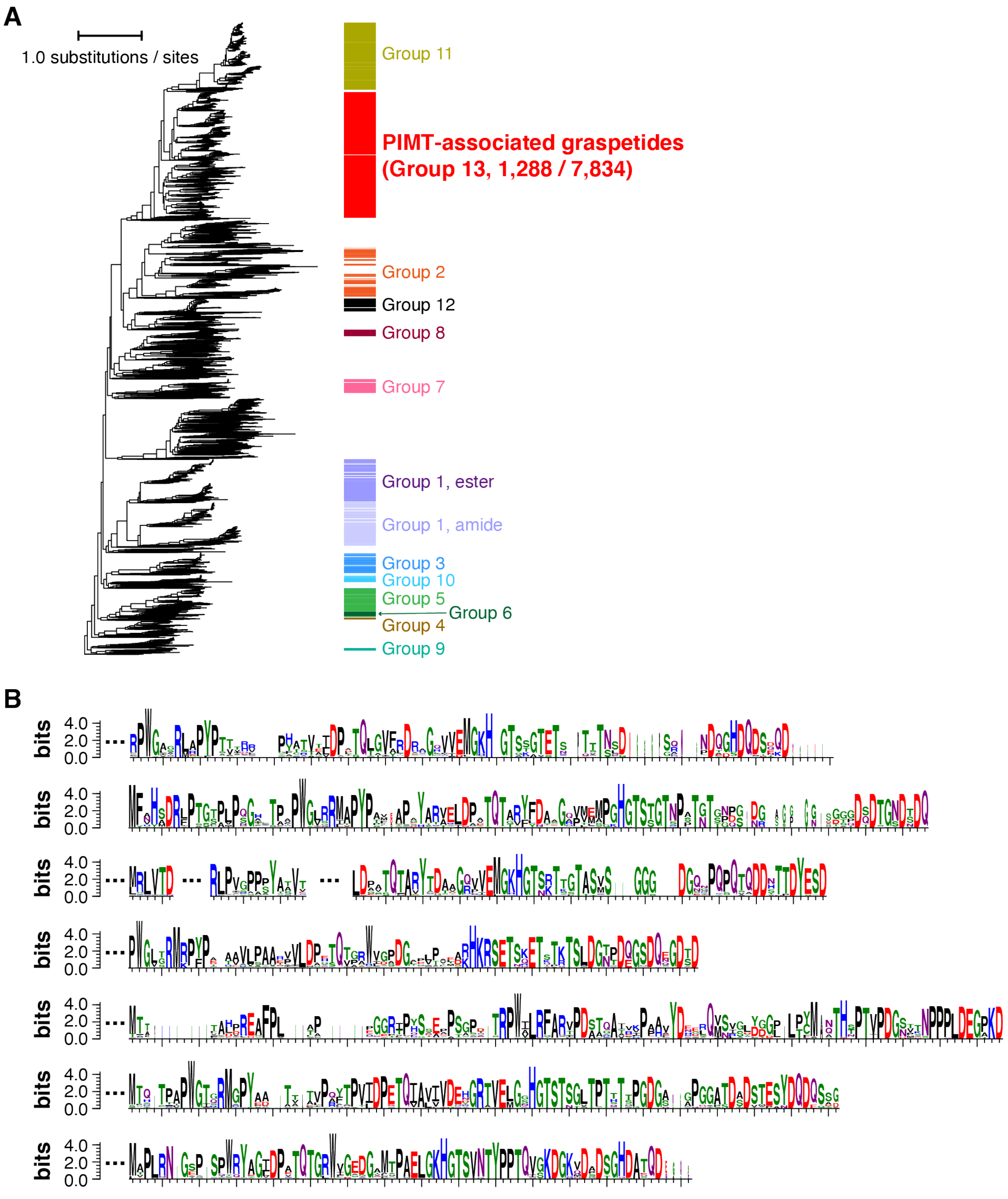


**Figure S1. Bioinformatic analysis of putative PIMT-modified graspetides.** (A) A maximum likelihood tree of 7,834 ATP-grasp enzymes obtained through PSI-BLAST. Previously reported ATP-grasp enzymes for group1-12 graspetides and enzymes for group 13 graspetides are annotated by color-coded strips. Lists of enzymes are provided in **Supplementary Dataset 2**. (B) Sequences of group 13 graspetides. Logos of precursors for seven subgroups whose members exceed 50 were generated by WebLogo.[20]

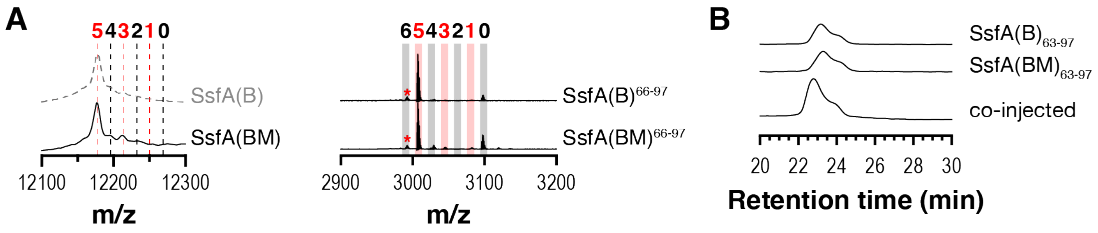


**Figure S2. MS and LC analysis of SsfA(B) and SsfA(BM).** (A) MALDI-TOF-MS spectrum of SsfA(BM) (left) and trypsin-cleaved SsfA(B) and SsfA(BM) (right). Mass spectrum of SsfA(B) (in Figure 2B) is shown in dotted line for comparison. Number of eliminated water molecules are designated above spectra. Red asterisks indicate the mass peaks resulting from the laser-induced deamination, which are not relevant to the activity of SsfB or SsfM. Observed and calculated mass value can be found in **Supplementary Dataset 3**. (B) HPLC analysis of SsfA(B)$_{63–97}$ and SsfA(BM)$_{63–97}$.

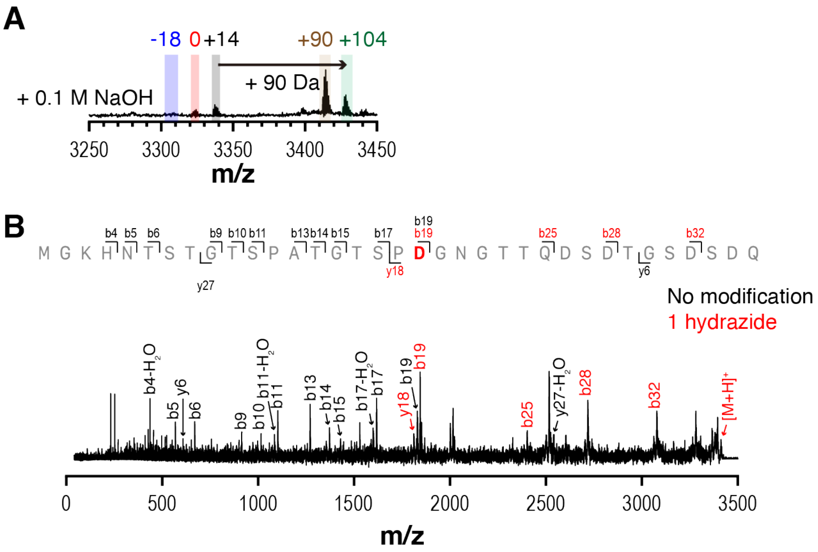


**Figure S3. Mass analysis of ester-hydrolyzed hydrazide-containing SsfA(BM)$_{63–97}$.** MALDI-TOF-MS spectrum (A) and MALDI-TOF/TOF-MS spectrum (B) were obtained to locate the hydrazide. Peptide was dissolved in 0.1 M NaOH solution and incubated at 25 $^{o}$C for 30 minutes. Relative mass value changes to SsfA(BM)$_{63–97}$ are given above each peak. Ions are colored according to the numbers of hydrazide group in the fragments (0, black; 1, red). Observed and calculated mass values can be found in **Supplementary Dataset 3**.

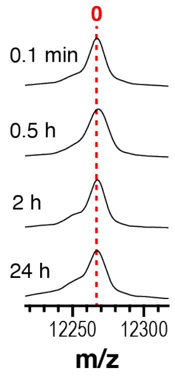


**Figure S4. MALDI-TOF-MS spectra of SsfM-mediated modification reaction of SsfA.** Substrate was incubated with SsfM at a same condition as in Figure 3B. Relative mass value changes to the unreacted substrates are given above each peak. Observed and calculated mass values can be found in **Supplementary Dataset 3**.

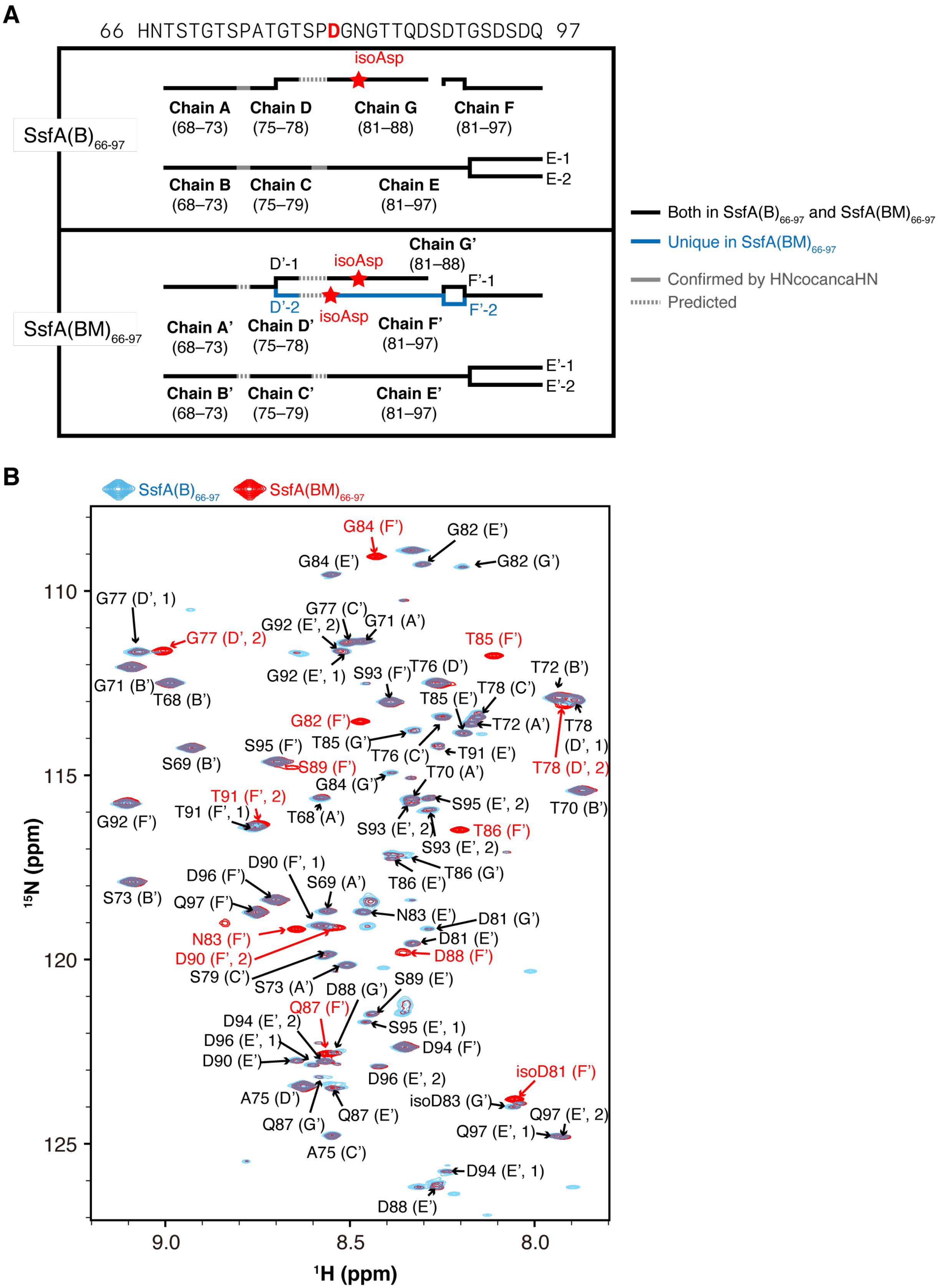


**Figure S5. Assigned Chains and $^1H$-$^{15}N$ HSQC spectra of SsfA(B)$_{66–97}$ and SsfA(BM)$_{66–97}$.** (A) Summary of the NMR-assigned chains. Signals that are observed in both SsfA(B)$_{66–97}$ and SsfA(BM)$_{66–97}$ spectra are annotated in black, while those exclusively observed in the spectrum of SsfA(BM)$_{66–97}$ are colored in blue. Observed isoaspartates are annotated (See Figures 3E, S6, and S7 for the detail). Gray solid lines indicate the connectivity confirmed by hNcocancaNH and gray dashed lines denote the predicted connectivity between peptide chains. (B) Overlaid $^{1}$H-$^{15}$N HSQC spectra of SsfA(B)$_{66–97}$ (blue) and SsfA(BM)$_{66–97}$ (red). Signals from the spectrum of SsfA(BM)$_{66–97}$ are labeled, with color indicating its presence in the spectrum of SsfA(B)$_{66–97}$ (black, observed in both spectra; red, exclusively present in SsfA(BM)$_{66–97}$). Chemical shift values can be found in **Supplementary Dataset 3**.

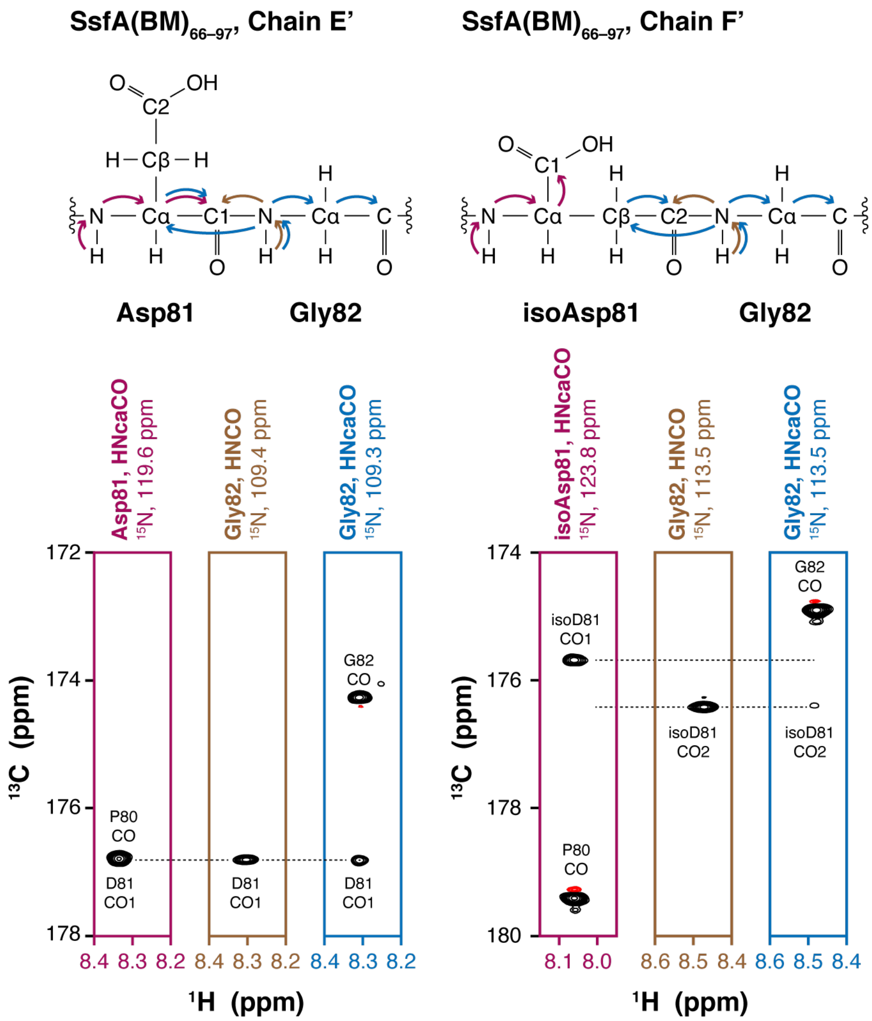


**Figure S6. HNcaCO and HNCO analysis of SsfA(BM)$_{66–97}$ chain E'/F'.** Magnetization transfer in Asp-Gly and isoAsp-Gly for HNcaCO/HNCO experiments (top) and HNcaCO/HNCO strip plots indicating the presence of aspartate and isoaspartate (bottom). Chemical shift values can be found in **Supplementary Dataset 3**.

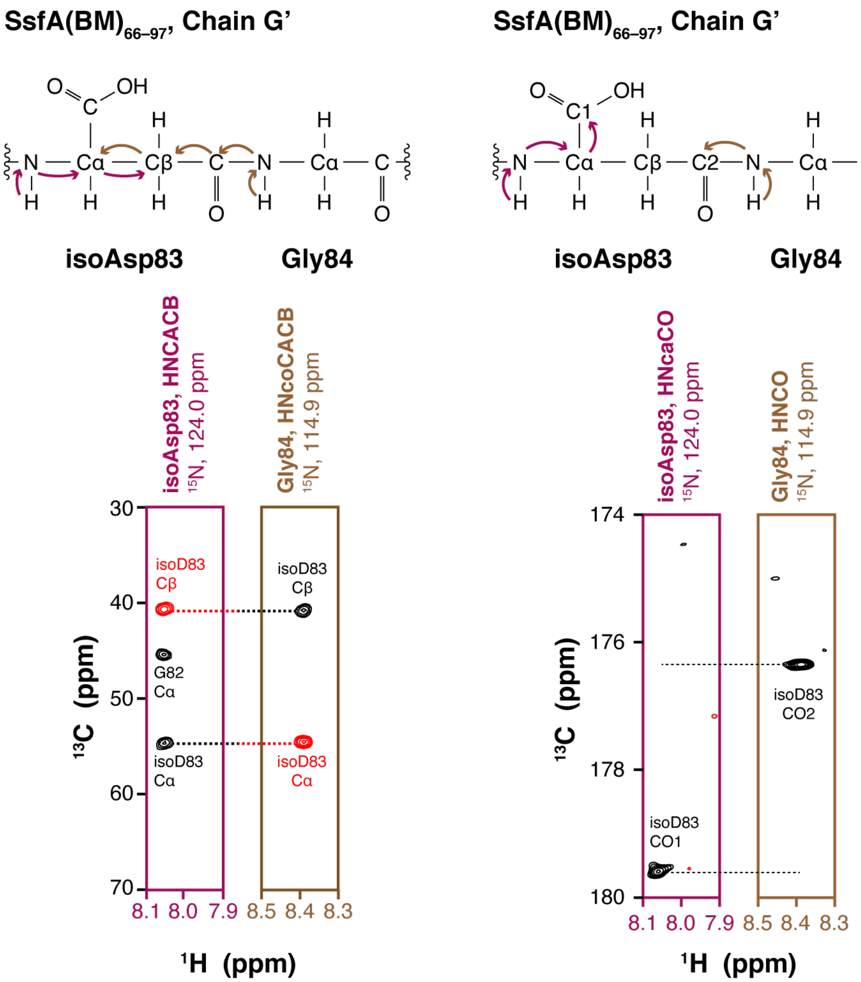


**Figure S7. HNCACB, HNcoCACB, HNCO, and HNcaCO analysis of SsfA(BM)$_{66–97}$ chain G'.** Magnetization transfer in isoAsp-Gly for HNCACB, HNcoCACB, HNcaCO, and HNCO experiments (top) and strip plots taken from the $^{1}H^{N}$-$^{13}C$ planes indicating the presence of isoaspartate (bottom). Chemical shift values can be found in **Supplementary Dataset 3**.

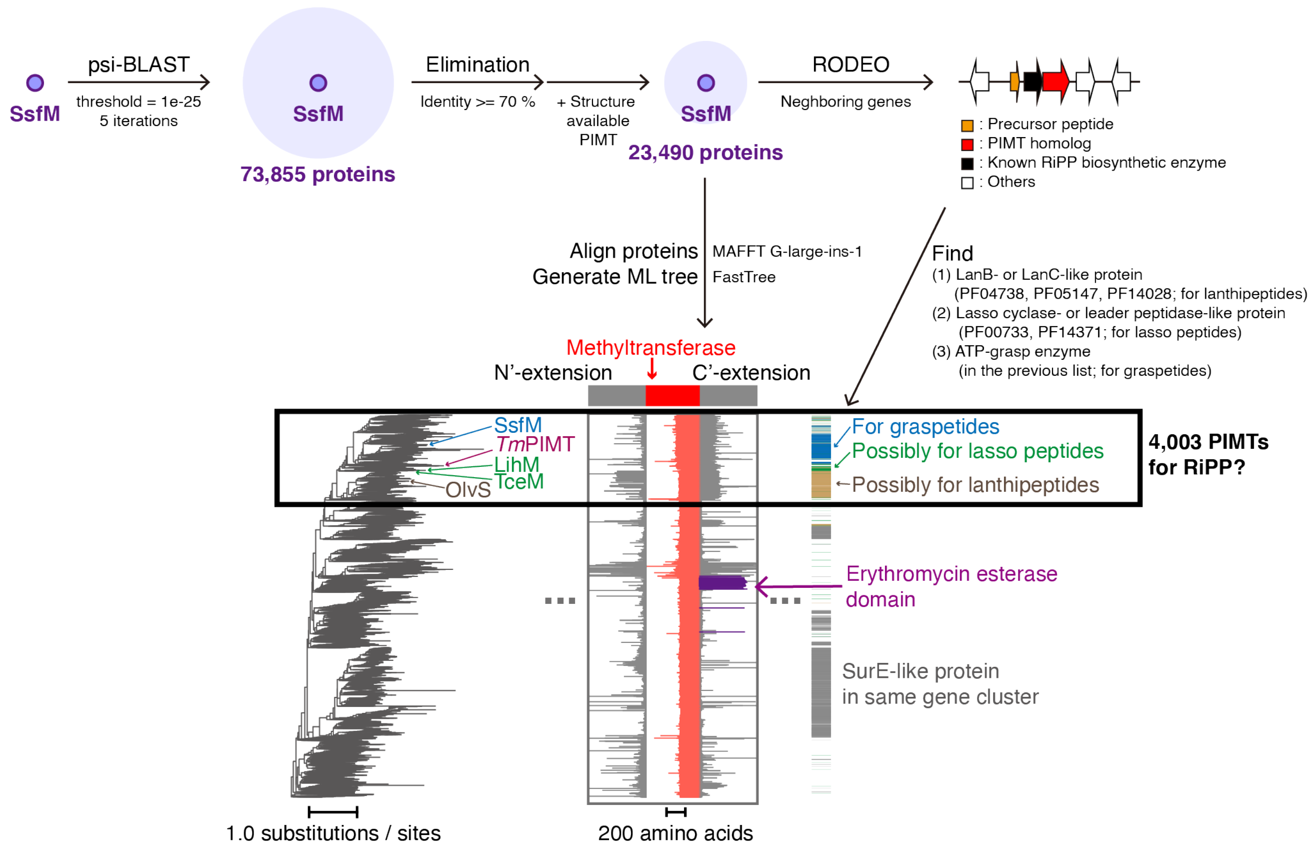


**Figure S8. Bioinformatic analysis workflow of PIMTs.** SsfM homologs were aligned and visualized in a maximum likelihood tree. Protein architectures and putative biological pathway of proteins are illustrated together. See **Materials and Methods** section for the details.

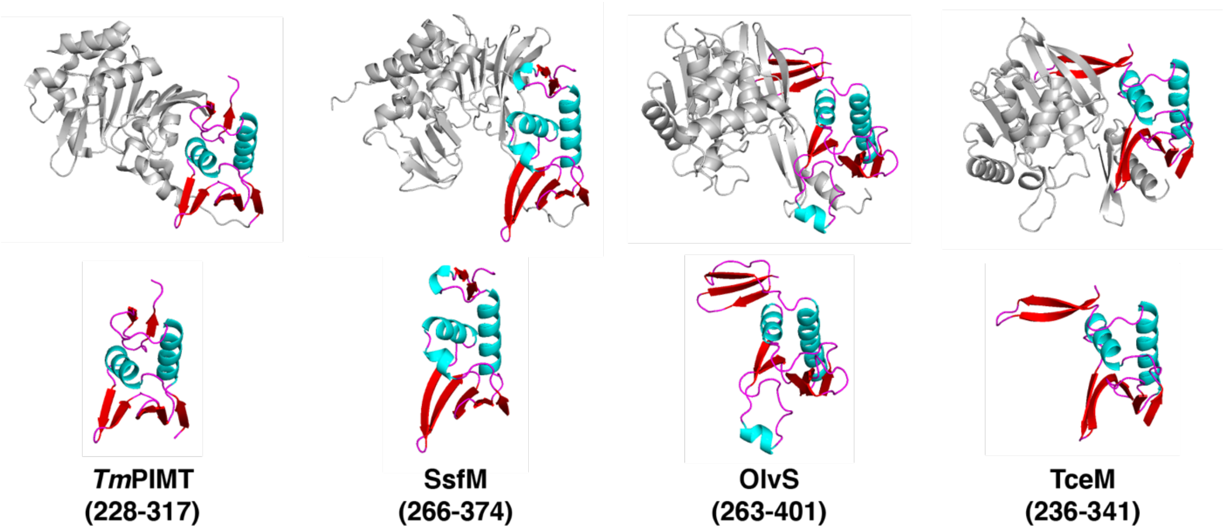


| | *Tm*PIMT | SsfM | OlvS | TceM |
|---|---|---|---|---|
| *Tm*PIMT | | | | |
| SsfM | 4.5 (2.8) | | | |
| OlvS | 3.6 (3.7) | 4.8 (3.5) | | |
| TceM | 5.7 (3.1) | 3.4 (4.4) | 7.3 (3.4) | |

**Z-score (RMSD, Å)**

**Figure S9. Structural analysis of extra C-terminal domains in model PIMTs.** Crystal structure of PIMT from *Thermotoga maritima* (PDB 1DL5)[25] and Alphafold structures of SsfM, OlvS, and TceM were used for the analysis. Residue numbers of the C-terminal domains are given in parentheses. Proteins were aligned by their C-terminal domains. Methyltransferase domain and secondary structural elements in C-terminal domains were annotated by color (gray, methyltransferase; cyan, α helix; red, β sheet; magenta, loop). *Z*-score values and RMSD values are given below.

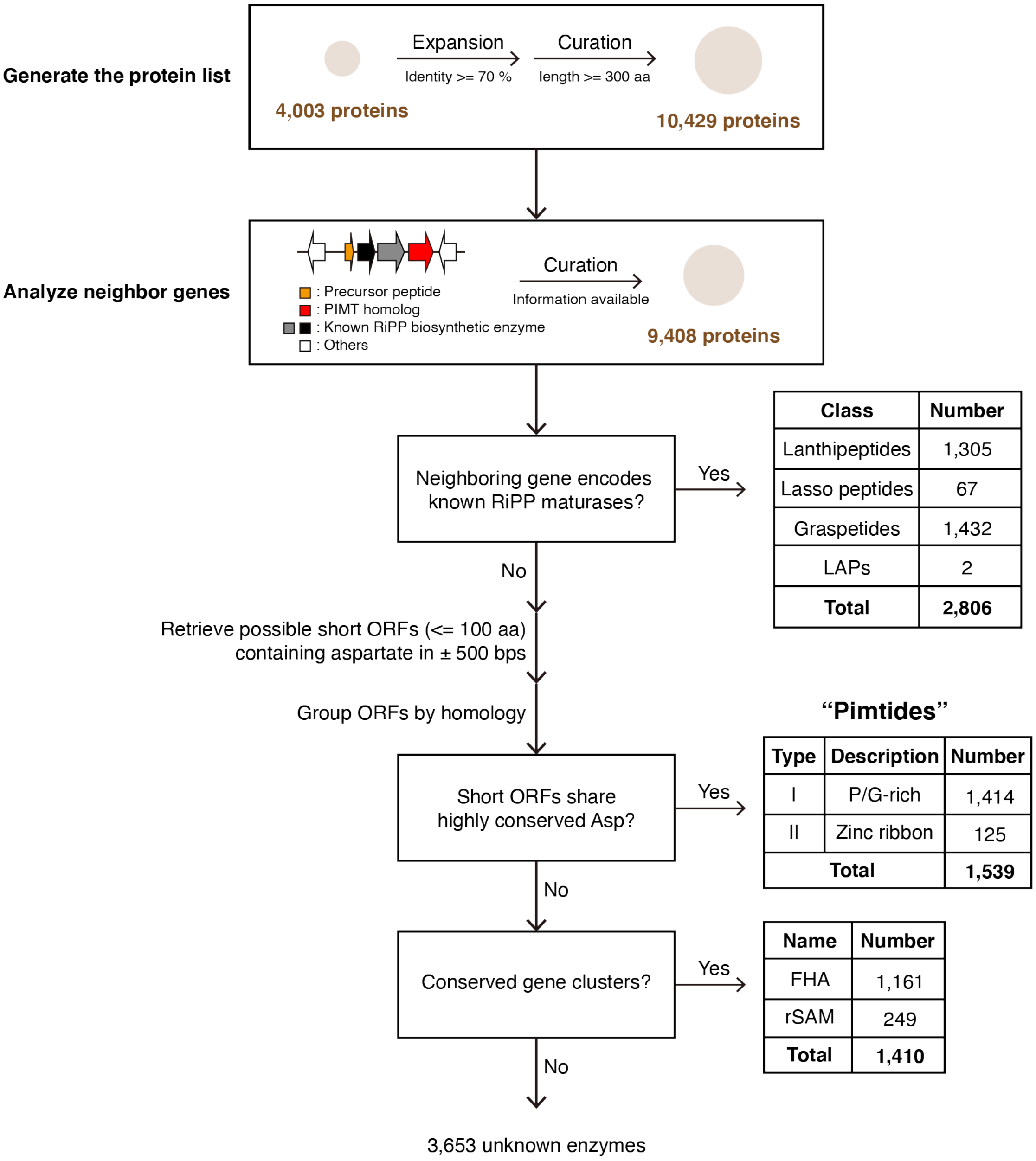


| Class | Number |
|---|---|
| Lanthipeptides | 1,305 |
| Lasso peptides | 67 |
| Graspetides | 1,432 |
| LAPs | 2 |
| **Total** | **2,806** |

**"Pimtides"**

| Type | Description | Number |
|---|---|---|
| I | P/G-rich | 1,414 |
| II | Zinc ribbon | 125 |
| **Total** | | **1,539** |

| Name | Number |
|---|---|
| FHA | 1,161 |
| rSAM | 249 |
| **Total** | **1,410** |

**Figure S10. Genome mining procedure of C-terminal domain-containing PIMTs.** See **Materials and Methods** section for the detail.

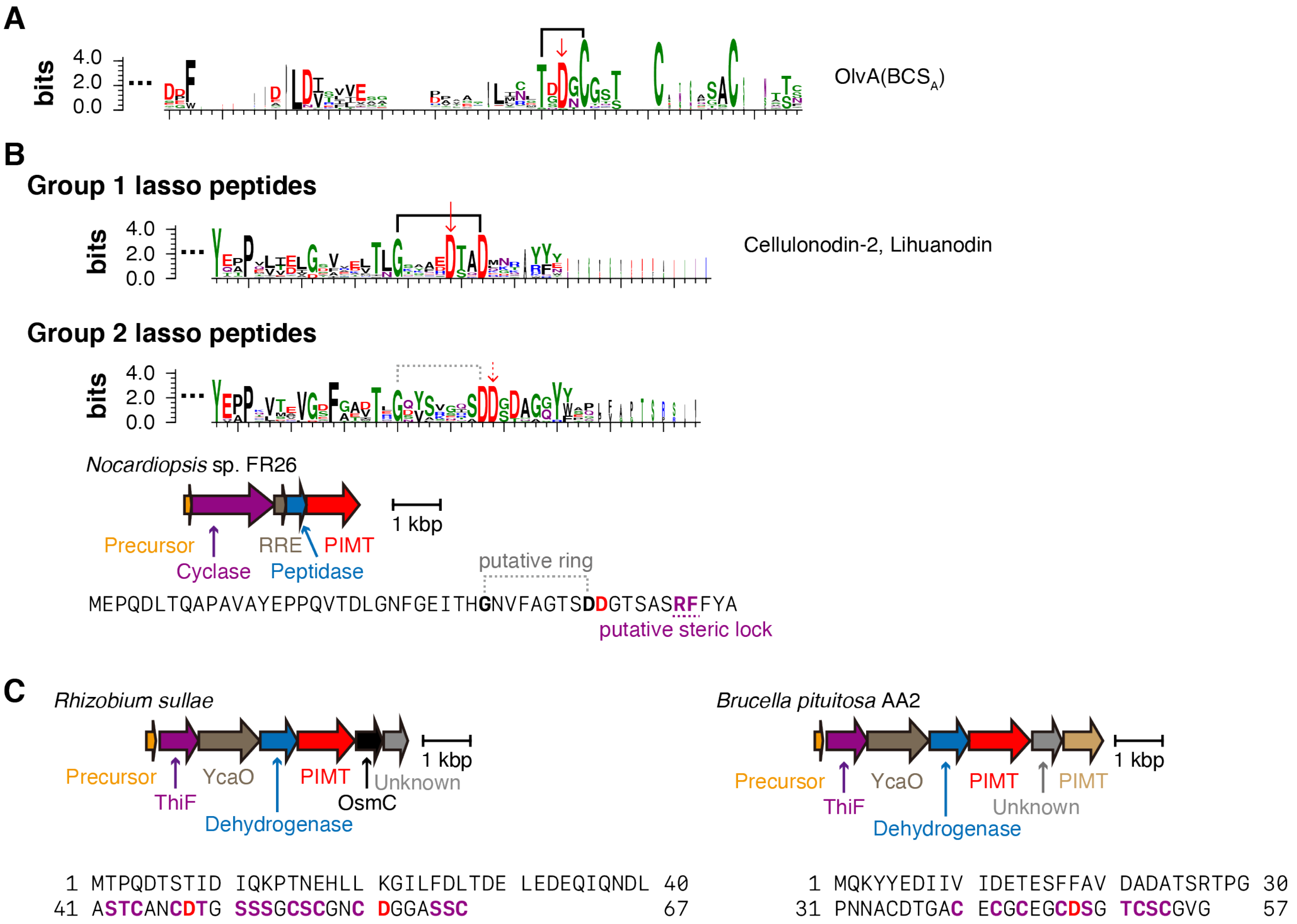


**Figure S11. Features of PIMT-associated RiPPs.** (A–B) Sequence logos of precursors for lanthipeptides (A) and lasso peptides (B). Previously identified thioether or isopeptide bond in lanthipeptides or group 1 lasso peptides are shown in black brackets, respectively. Predicted isopeptide bond in group 2 lasso peptides are drawn in gray dashed bracket. Modified aspartates in mature peptides are highlighted in red solid arrows, while predicted one in group 2 lasso peptide is highlighted in red dashed arrows. Biochemically characterized members of lanthipeptides and group 1 lasso peptides are annotated.[26,27] A model BGC and precursor for the group 2 lasso peptides are shown below. (C) Biosynthetic gene clusters of LAPs containing PIMT-encoding genes. Arrows are color-coded according to the conserved protein domain. Precursor peptide sequences are given below each BGCs. Putative modified aspartates in each precursor are highlighted in red. Another PIMT that lacks the C-terminal domain (brown) is encoded in the same BGC from *B. pituitosa*.

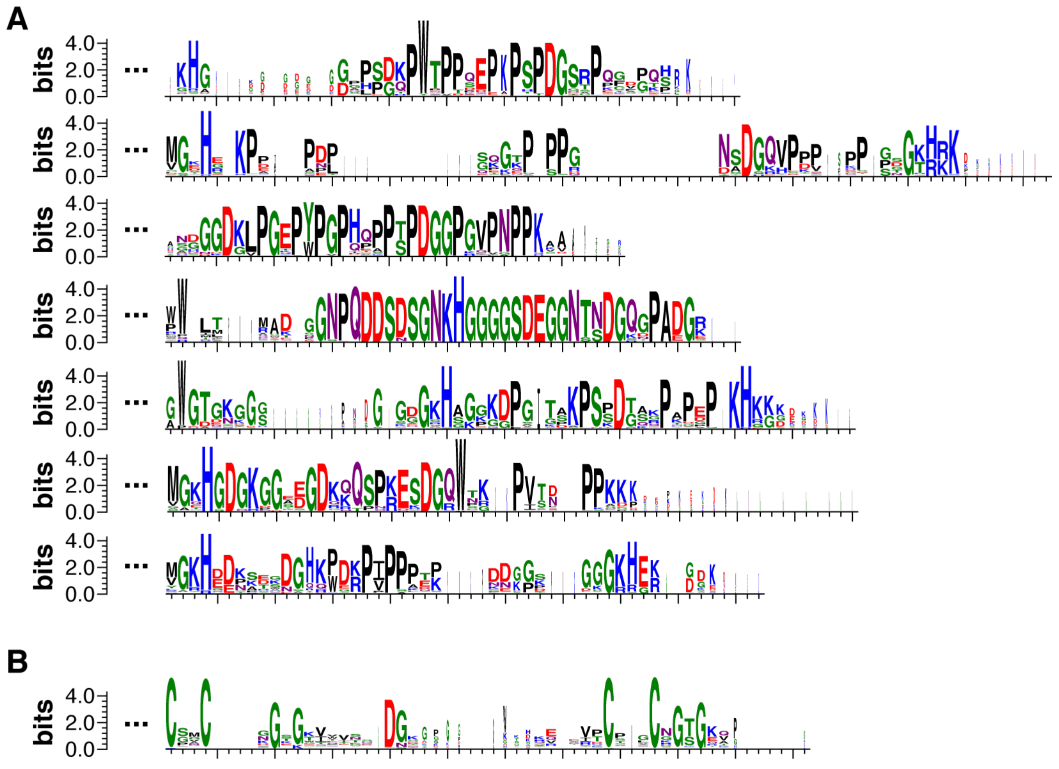

**Figure S12. Sequence logos of precursor peptides for pimtides.** Logos of type I precursors for seven subgroups whose members exceed 50 (A) and type II precursors (B) were generated by WebLogo.[20]

**A**

| | | | |
|---|---|---|---|
| ***Ec*DnaJ** | 152 | ..AKPGTQPQTCPTCHGSGQVQMRQGFFAVQQTCPHCQGRGTLIKDP.. | 186 |
| **FcaA** | 1 | MGKHAKPVACPTCNGSGKITVTSDGKNETVSCGVCKGSGKA | 41 |
| **Consensus** | | P CPTC GSG C C G G | |

**B**

**Zinc ribbon of *Ec*DnaJ (PDB 1EXK)**

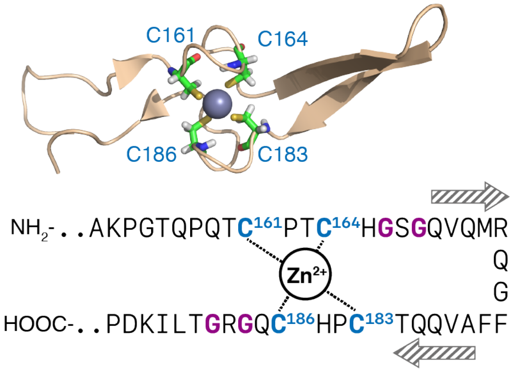


**Figure S13. Structure of zinc ribbon.** (A) Sequence alignment of zinc ribbon from *E. coli* DnaJ (*Ec*DnaJ) and FcaA. Commonly observed residues are shown below, and conserved CxxCxGxG motifs are highlighted (C, blue; G, purple). (B) Structure of the zinc ribbon of *Ec*DNAJ (PDB 1EXK).[28] Observed secondary structural elements and metal-ligand interactions are shown at the right.

**A**

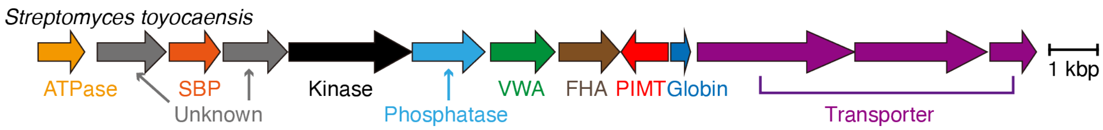


**FHA**

```
001 MPTCPNGHQSGSDDWCEVCGHRMAGAVPPPPPPPPAPGGGYGFPPPGGPGGPGGPGGYGGPGGYGGPGGPGGPGGPGGRT 080
081 ELCPQCRTPREGGAPFCEECRWNFLTNTATSYTPAAPRPPAPGPGPGGPFQQQPPGPSYGGDSFDYQGSRPSQMNRPAEP 160
161 IPPGPPGFGGDPSRPVPPPPGPGGHGGPGGHGGHGGPGAPQAFQPGPPAPPGFPQDTGRPQQGGPSFGGGDDDWVISPPG 240
241 GPGGPGGPGGPNGPGQGGGYGYPQPGGTQAPPGPGGFPQQPRPPQGPATWTATIGPDREYFMAMMQRSGPEAAGLNLPAY 320
321 SPEQQRQLTGNQITIGRRRHSTGDTPDIDLSTPPEDPGVSHQHAVLVQQPDGTWAVVDQNSTNGTTVNGSDEPIQPFVPV 400
401 PLQDGDRVHVGAWTTITIRRG 421
```

**B**

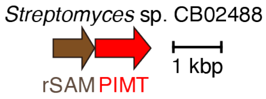


**Figure S14. Conserved PIMT-associated gene clusters.** Representative gene clusters of FHA/VWA-associated PIMT (A) and rSAM-associated PIMT (B) are shown. Arrows are color-coded according to the predicted protein domain. Sequence of an FHA domain-containing protein is shown below with colors illustrating the P/G-rich region (blue) and aspartates in the region (red).

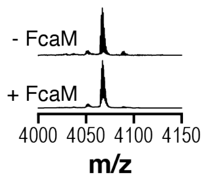


**Figure S15. MALDI-TOF-MS spectra of the purified recombinant FcaA and FcaA(M).** Calculated and observed mass values can be found in **Supplementary Dataset 3**.

**Figure S16. NMR spectra of FcaA(M)$_{19–26}$.** FcaA(M)$_{19–26}$ was analyzed by $^1$H (A), $^1$H-$^1$H COSY (B), $^1$H-$^1$H TOCSY (C), and $^1$H-$^1$H NOESY (D). Chemical shift values can be found in **Supplementary Dataset 3**.

**A**

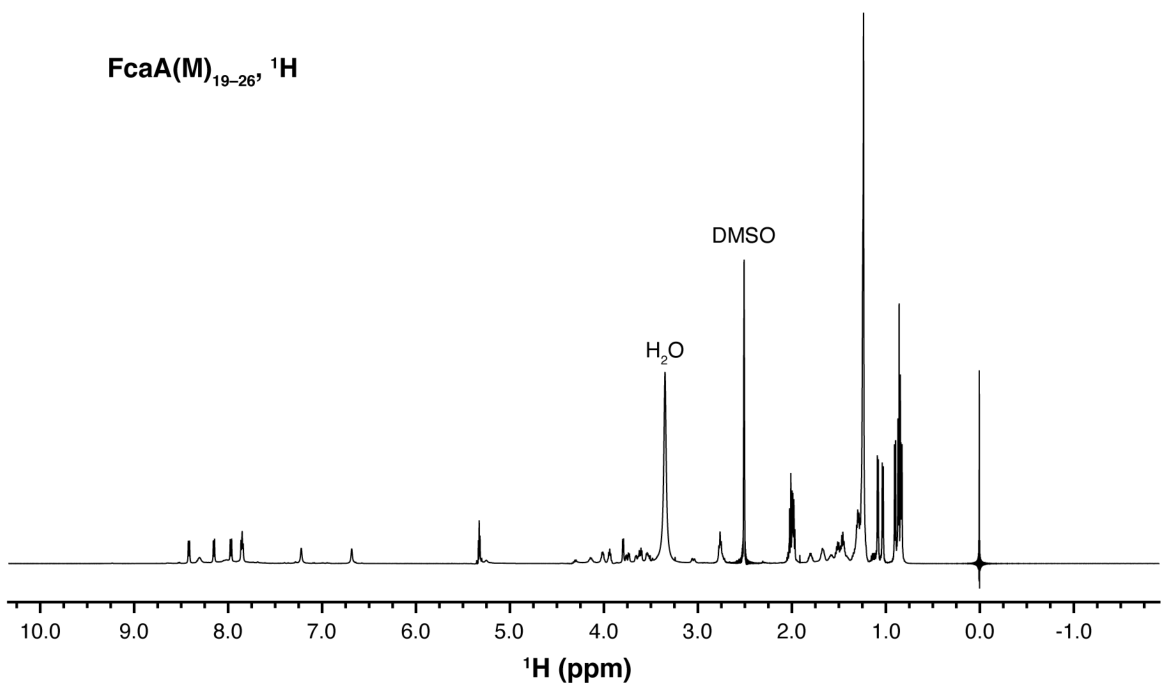


**B**

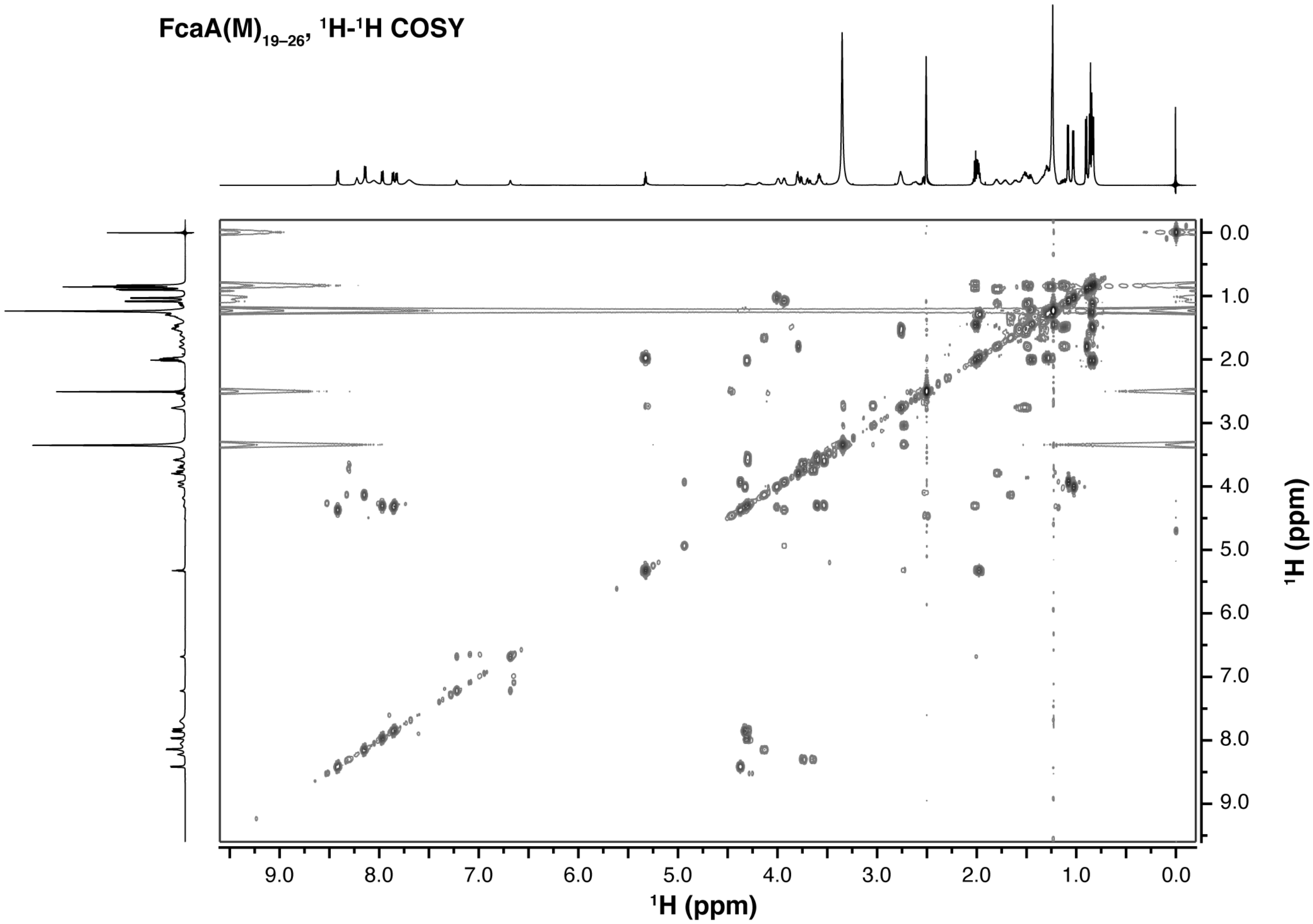

**C**

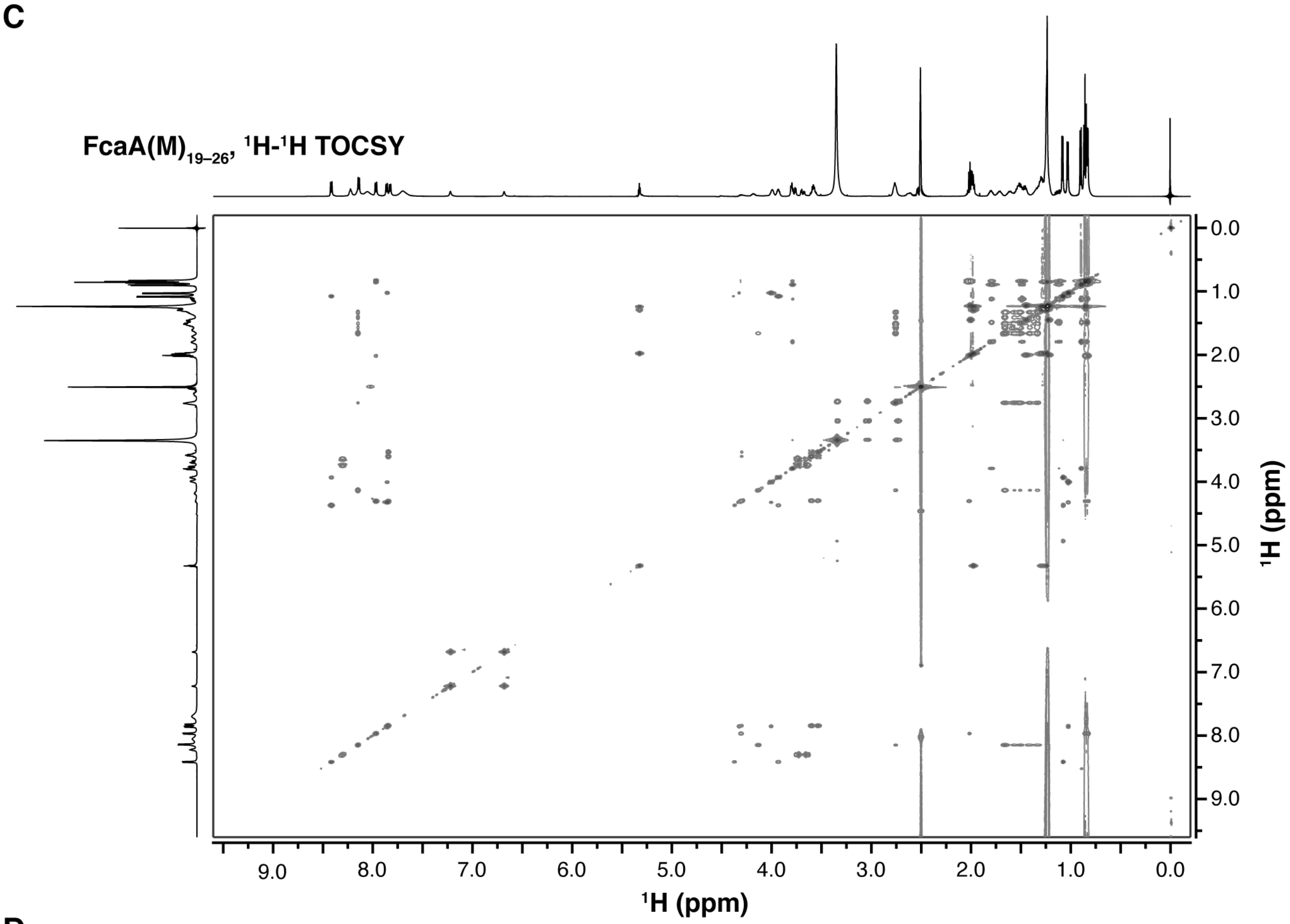
FcaA(M)$_{19–26}$, $^1$H-$^1$H TOCSY
9.0 8.0 7.0 6.0 5.0 4.0 3.0 2.0 1.0 0.0
$^1$H (ppm)
0.0 1.0 2.0 3.0 4.0 5.0 6.0 7.0 8.0 9.0
$^1$H (ppm)

**D**

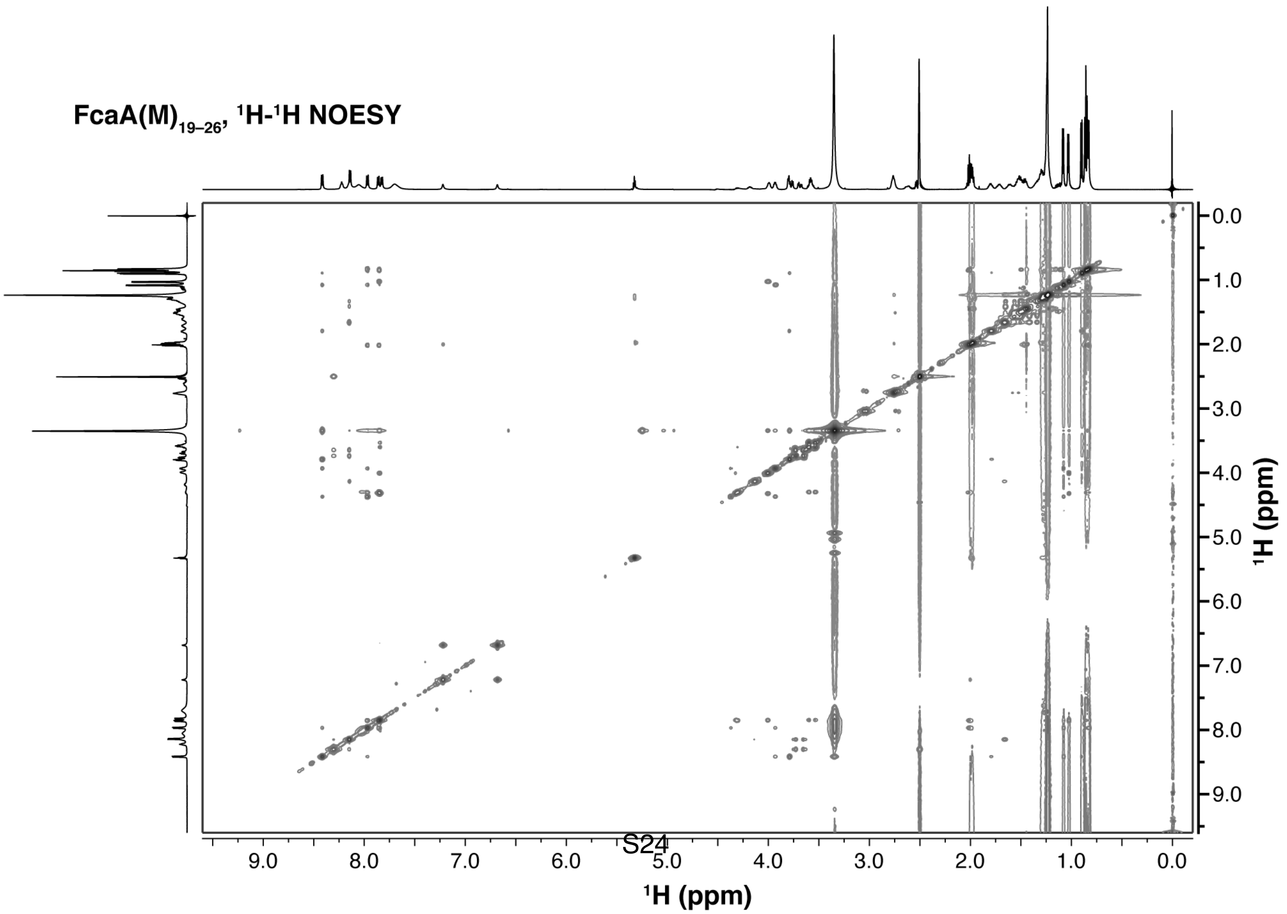
FcaA(M)$_{19–26}$, $^1$H-$^1$H NOESY
9.0 8.0 7.0 6.0 5.0 4.0 3.0 2.0 1.0 0.0
$^1$H (ppm)
0.0 1.0 2.0 3.0 4.0 5.0 6.0 7.0 8.0 9.0
$^1$H (ppm)

**Figure S17. NMR analysis of synthetic ITVTSDGK and ITVTS(isoD)GK.** (A–B) TOCSY and NOESY spectra of ITVTSDGK (A) and ITVTS(isoD)GK (B). In ITVTSDGK, of the two possible NOE signals, one between D6 $H^{\alpha}$ and G7 $H^{N}$ (red solid circle; A) and the other between D6 $H^{\beta}$ and G7 $H^{N}$ (black dotted circle; A), the former signal was uniquely observed. On the other hand, an NOE signal was observed between isoD6 $H^{\beta}$ and G7 $H^{N}$ (red solid circle; B), instead of isoD6 $H^{\alpha}$ and G7 $H^{N}$ (black dotted circle). (C–J) Raw $^{1}H$ (C, ITVTSDGK; G, ITVTS(isoD)GK), $^{1}H$-$^{1}H$ COSY (D, ITVTSDGK; H, ITVTS(isoD)GK), $^{1}H$-$^{1}H$ TOCSY (E, ITVTSDGK; I, ITVTS(isoD)GK), and $^{1}H$-$^{1}H$ NOESY (F, ITVTSDGK; J, ITVTS(isoD)GK) spectra. Observed chemical shift values can be found in **Supplementary Dataset 3**.

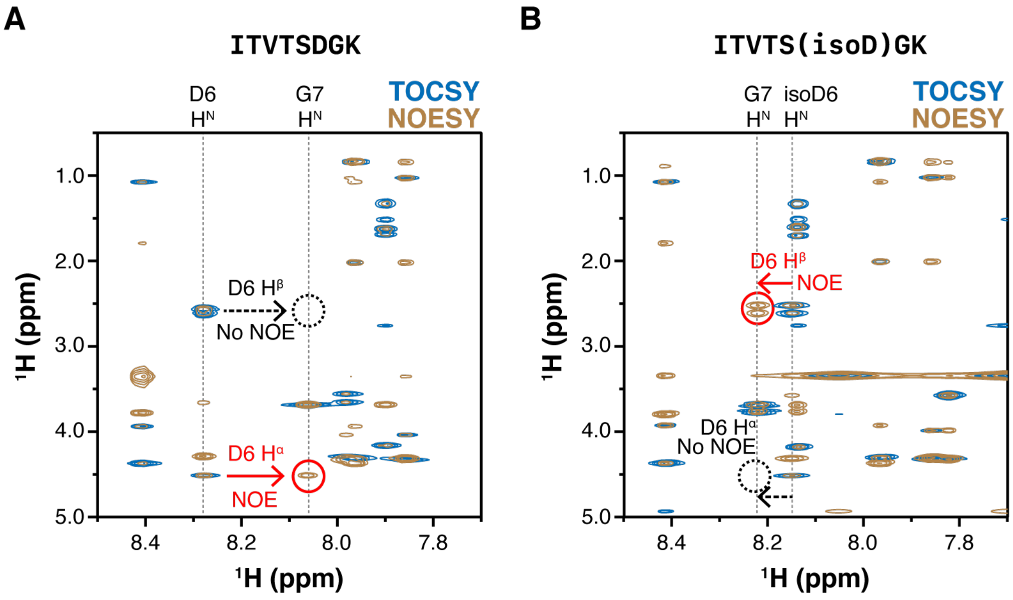

**C**

**ITVTSDGK, ¹H**

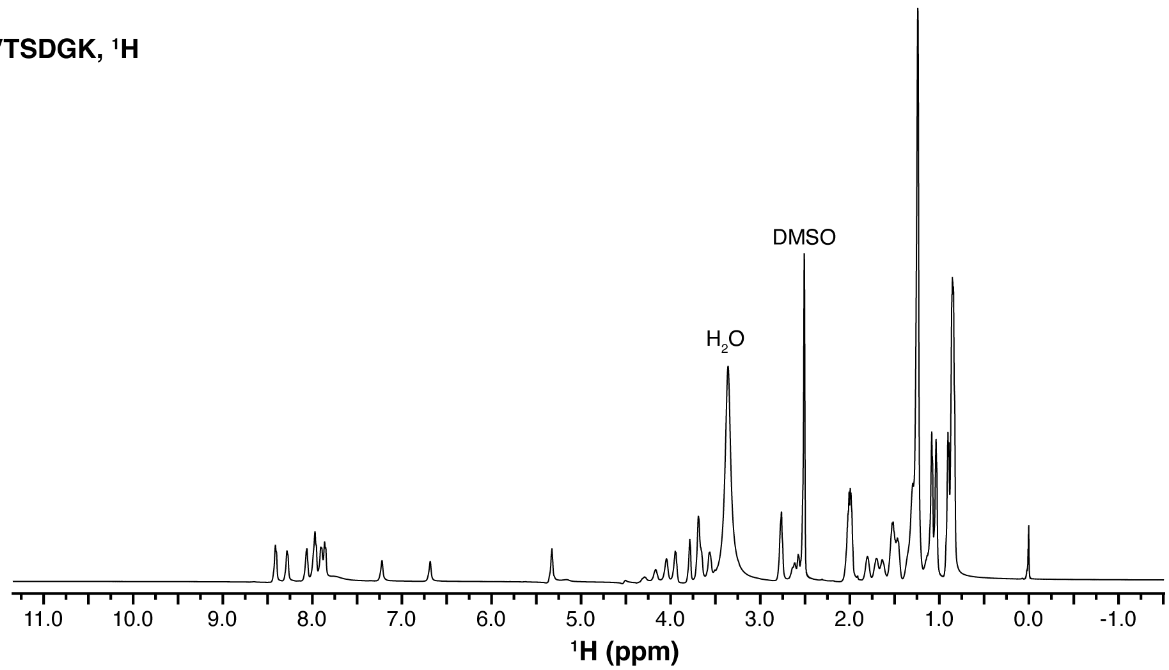


**D**

**ITVTSDGK, ¹H-¹H COSY**

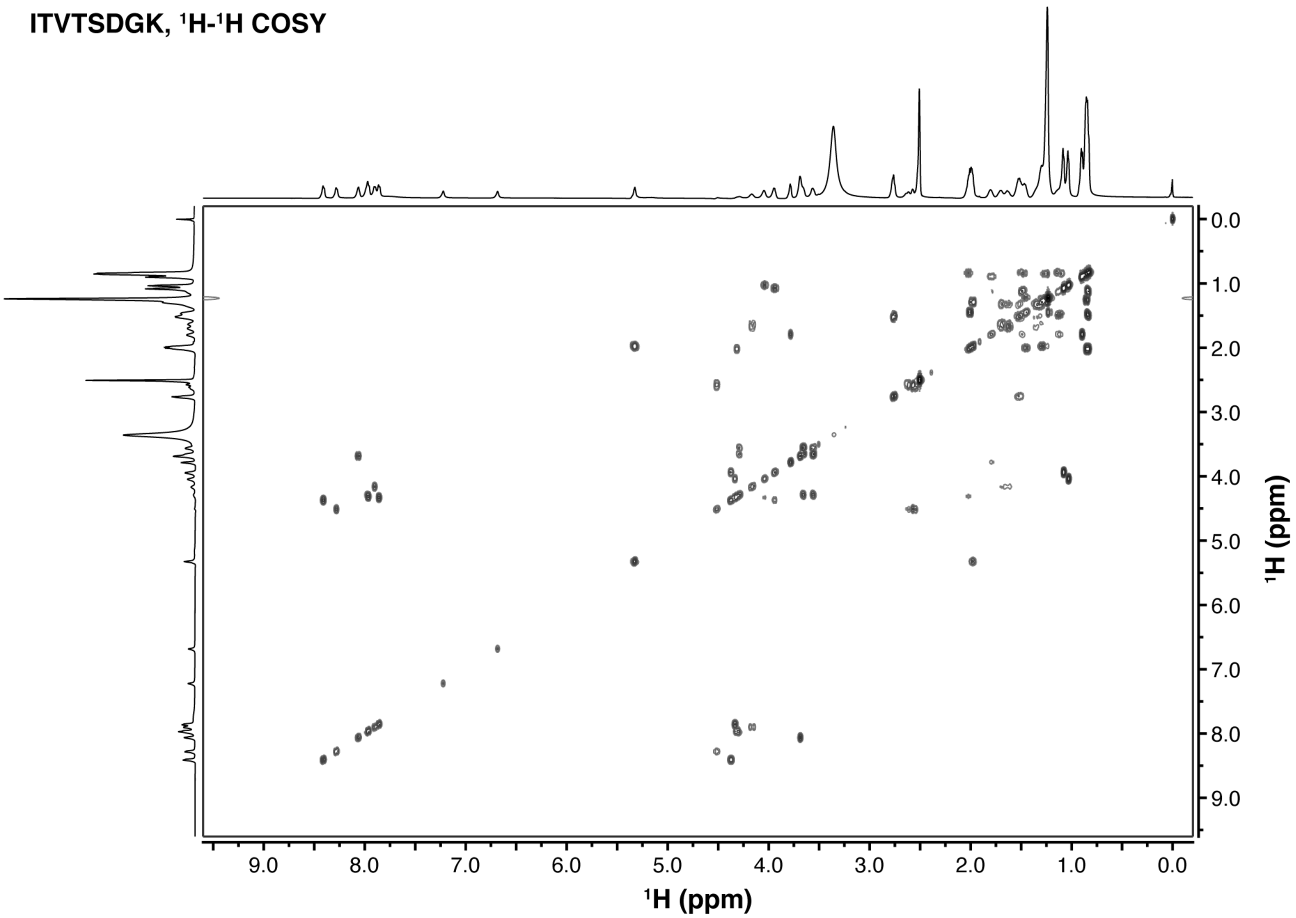

**E**

**ITVTSDGK, $^{1}H$-$^{1}H$ TOCSY**

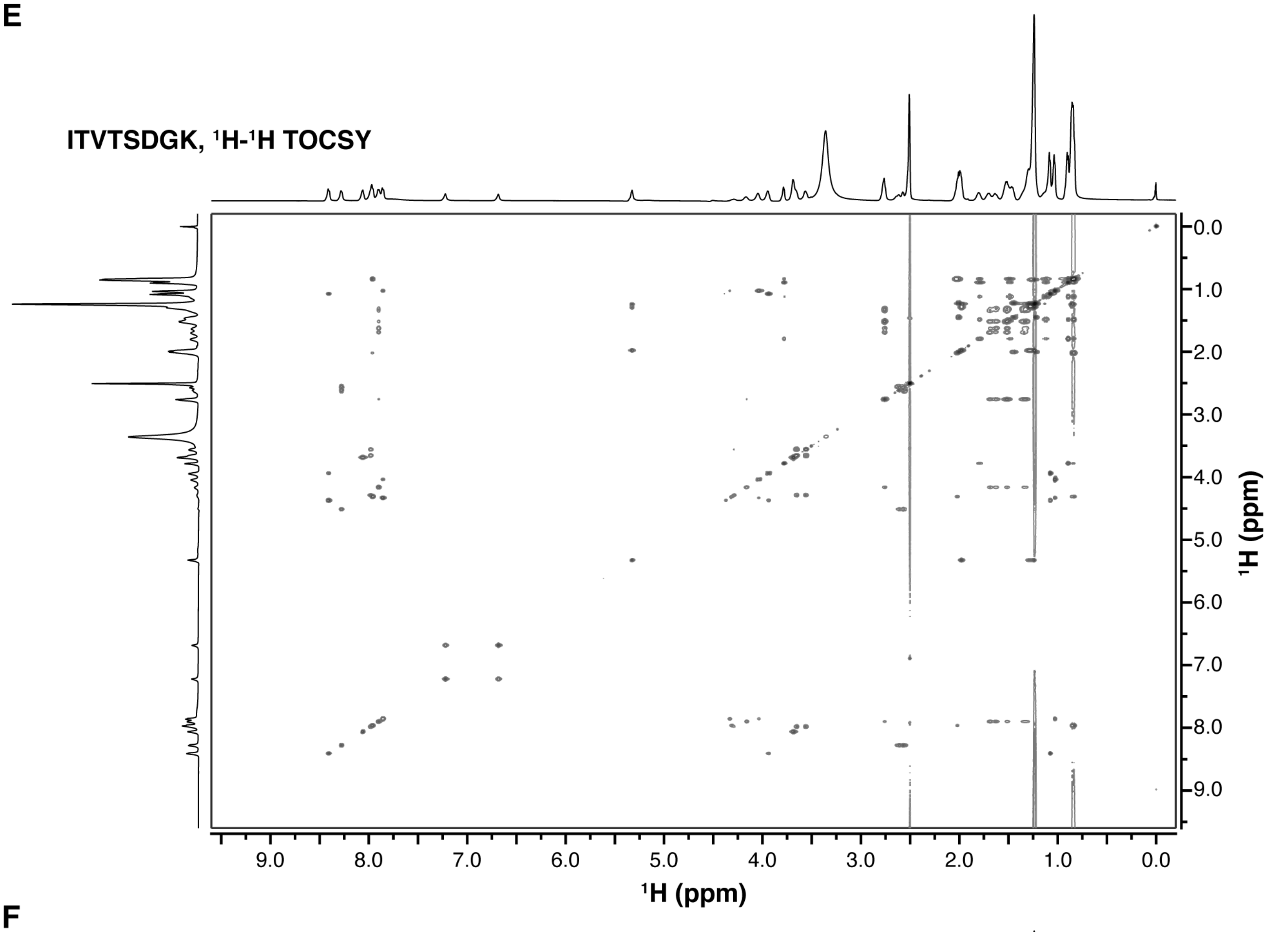


**F**

**ITVTSDGK, $^{1}H$-$^{1}H$ NOESY**

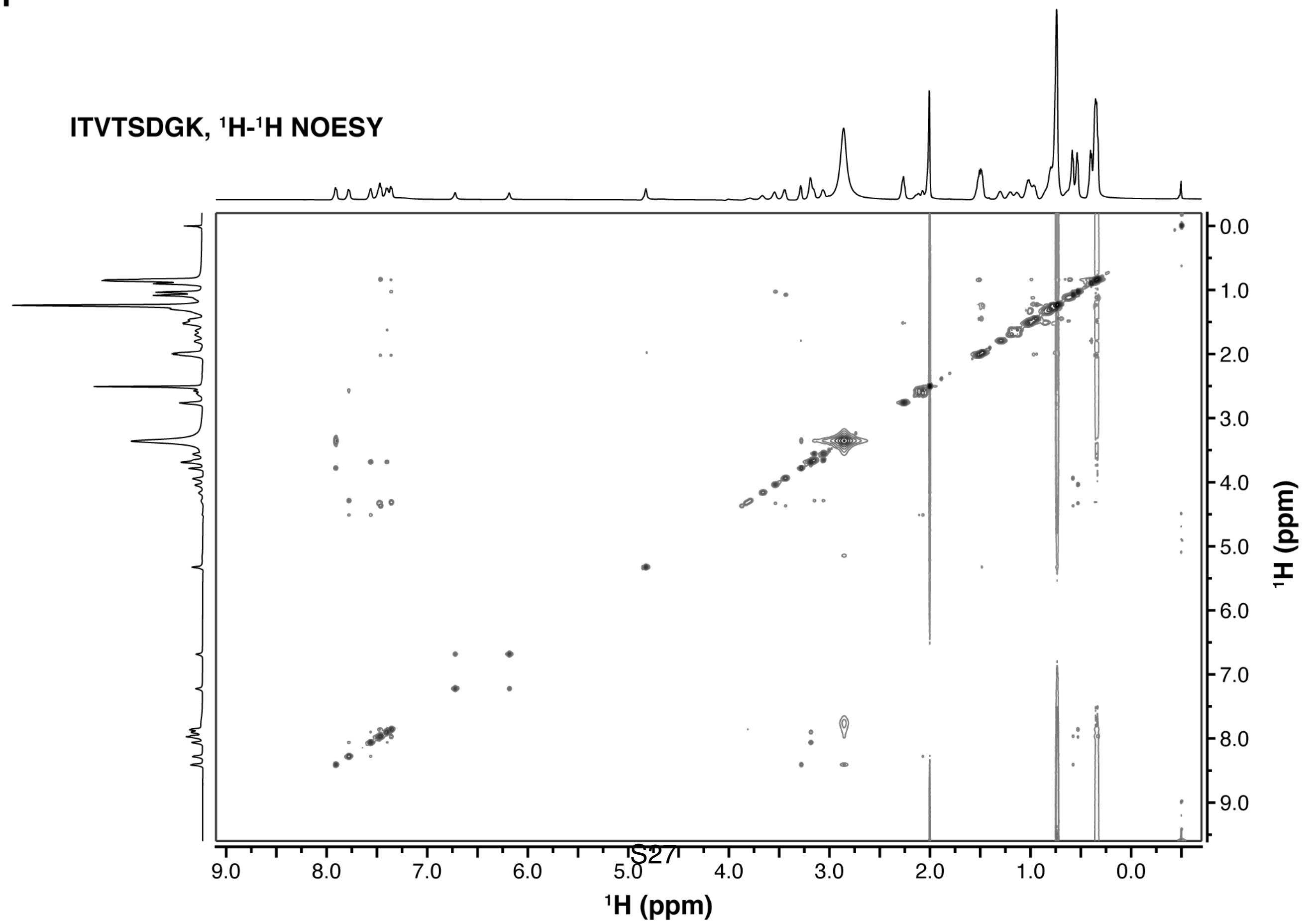

**G**

**ITVTS(isoD)GK, $^1H$**

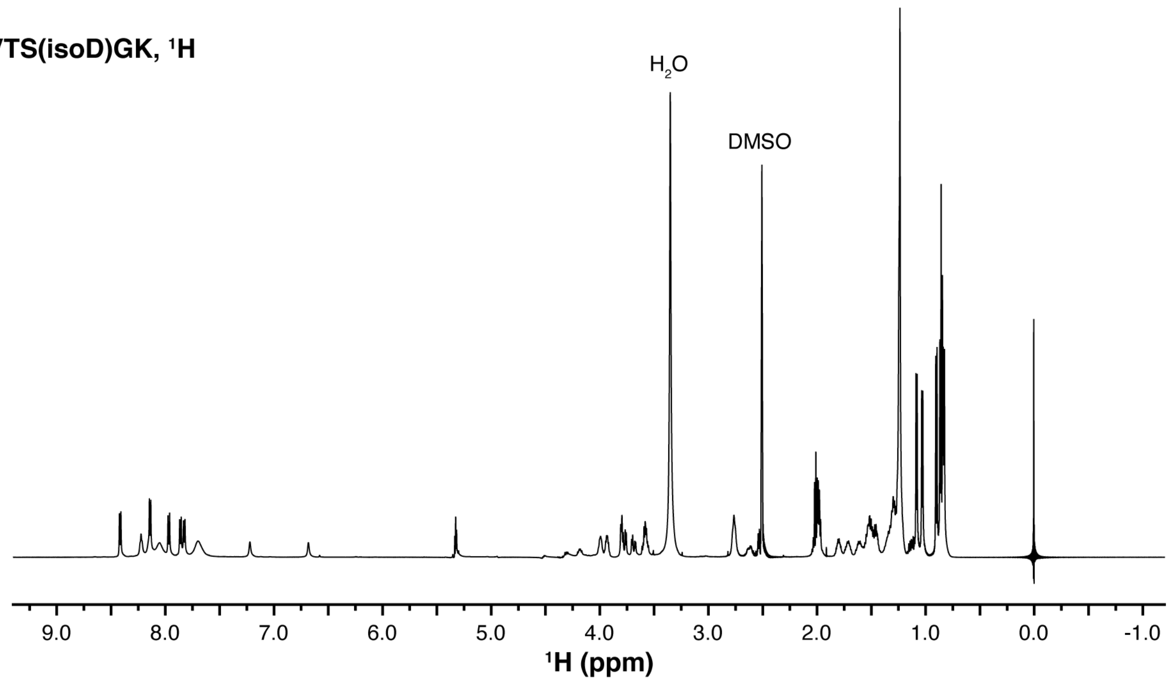


**H**

**ITVTS(isoD)GK, $^1H$-$^1H$ COSY**

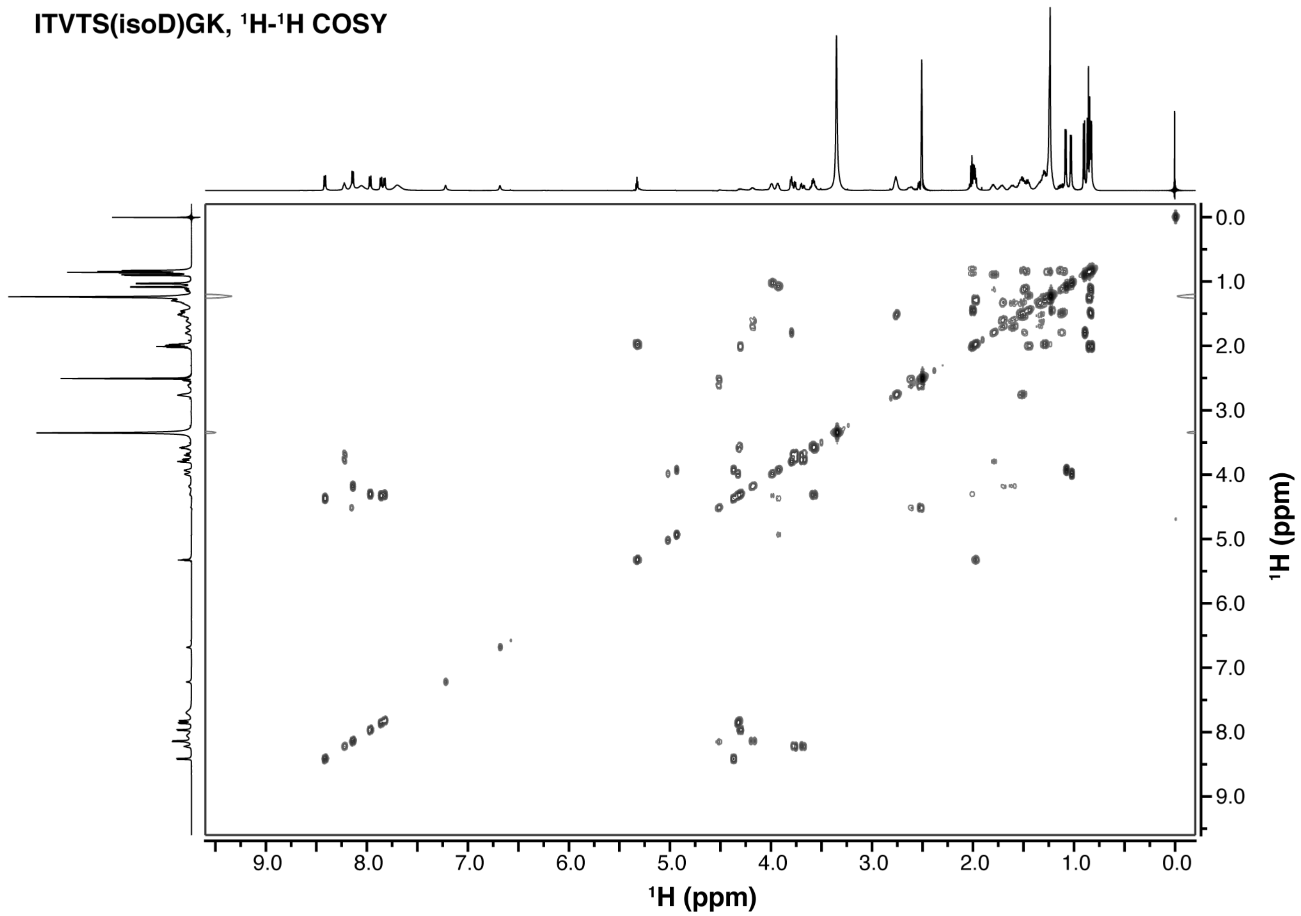

**I**

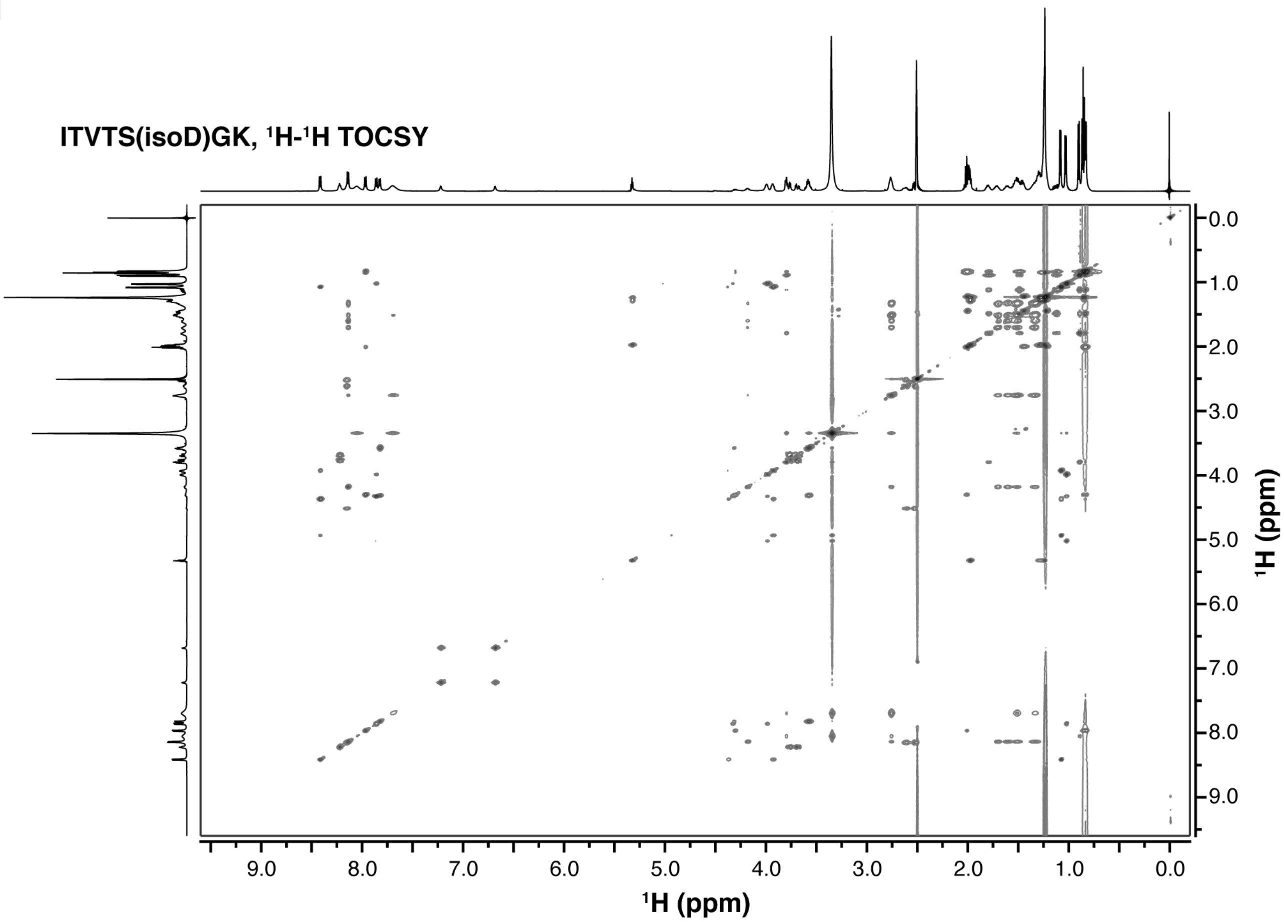
ITVTS(isoD)GK, $^1$H-$^1$H TOCSY
0.0
1.0
2.0
3.0
4.0
5.0
6.0
7.0
8.0
9.0
$^1$H (ppm)
9.0 8.0 7.0 6.0 5.0 4.0 3.0 2.0 1.0 0.0
$^1$H (ppm)

**J**

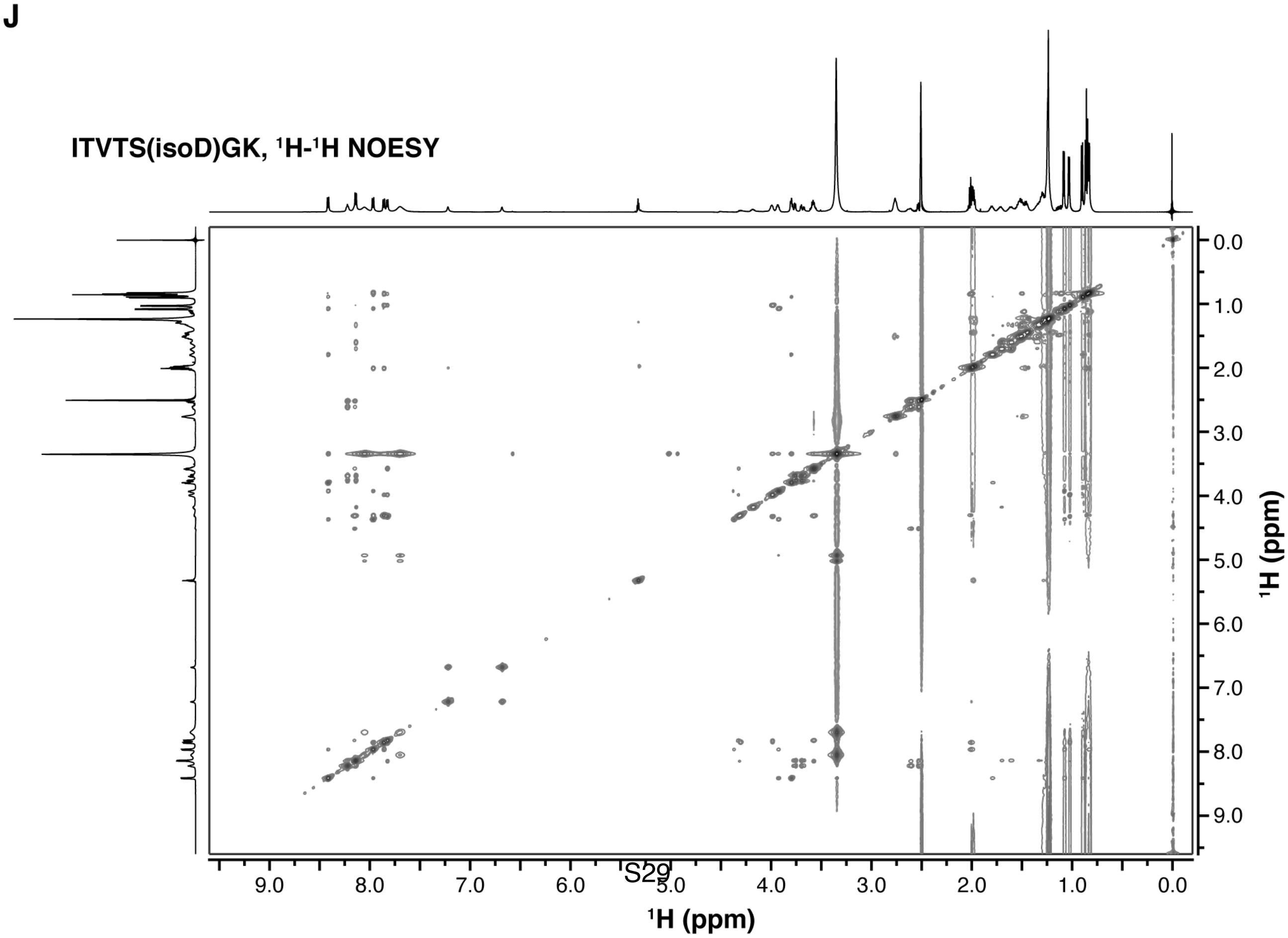
ITVTS(isoD)GK, $^1$H-$^1$H NOESY
0.0
1.0
2.0
3.0
4.0
5.0
6.0
7.0
8.0
9.0
$^1$H (ppm)
9.0 8.0 7.0 6.0 5.0 4.0 3.0 2.0 1.0 0.0
$^1$H (ppm)

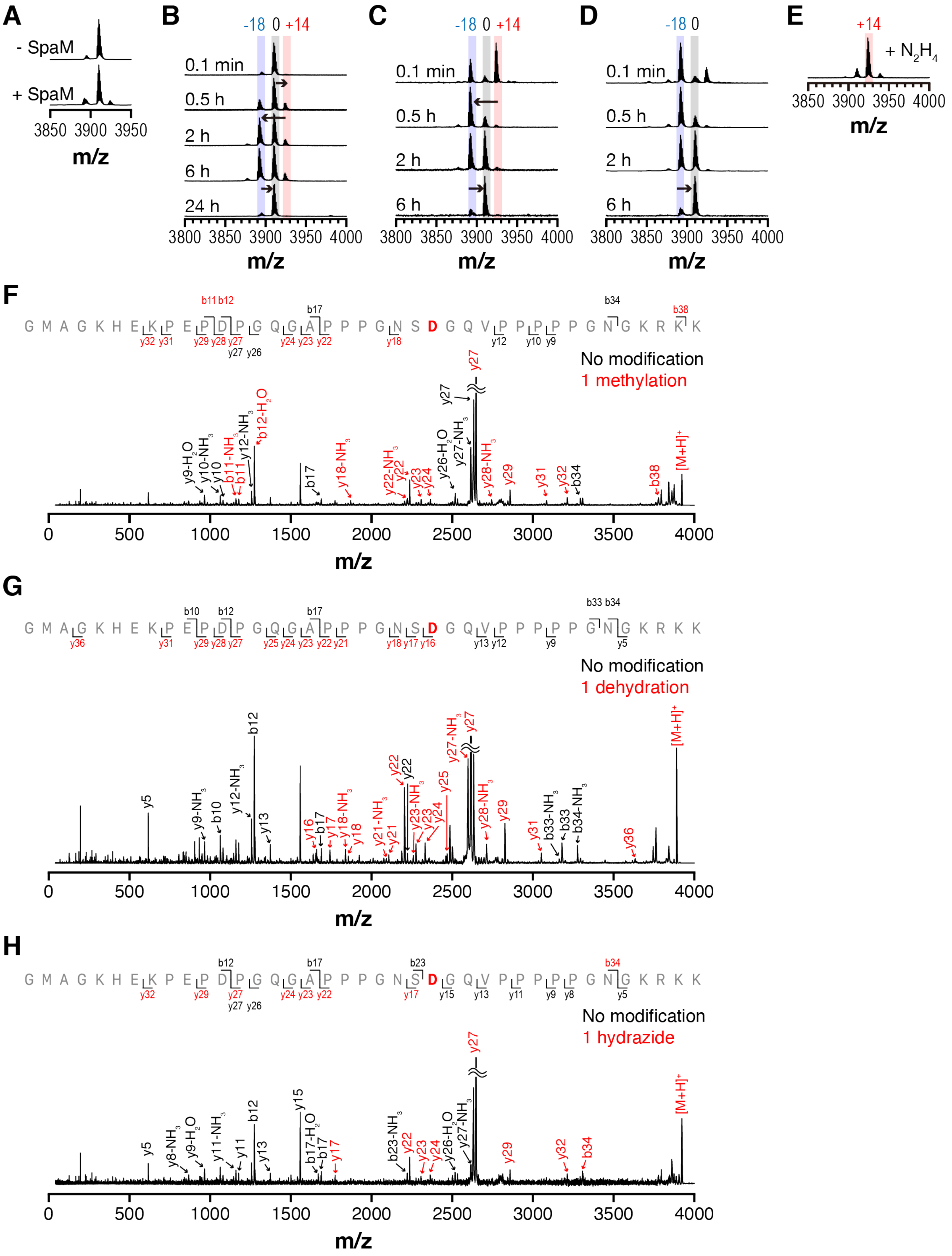


**Figure S18. Mass analysis of a precursor-enzyme for type I pimtide.** (A) MALDI-TOF-MS spectra of the purified recombinant SpaA (top) or SpaA(M) (bottom). (B) MALDI-TOF-MS spectra of SpaM-mediated SpaA modification *in vitro*. SpaA (20 µM) was mixed with SpaM (5 µM) in presence of DTT (1 mM), SAM (1 mM), and Tris-HCl pH 8.0 (50 mM) at 25 °C. Reaction was monitored at designated time points by mass analyzer. Relative mass value changes to the intact SpaA are given above peaks. (C–D) MALDI-TOF-MS spectra of spontaneous rearrangement of methylester- (D) or aspartimide-containing intermediates (E). Reaction conditions for Figure S6B–S6C were adopted to observe the spontaneous rearrangement of the intermediates. (E) MALDI-TOF-MS spectrum of hydrazide-containing peptide. Same reaction condition as Figure S6D was applied to the aspartimide-containing intermediate to generate hydrazide. Relative mass value to the intact SpaA is given above the peak. (F–H) MALDI-TOF/TOF-MS spectra of methylester- (F), aspartimide- (G), or hydrazide-containing SpaA (H). Ions are colored according to the numbers of modified residues in the fragments (0, black; 1, red). Calculated and observed mass values can be found in **Supplementary Dataset 3**.

### Tables

**Table S1. Optimized sequences of precursor peptides, ATP-grasp enzymes, and PIMTs for heterologous expression in *E. coli*.** Vectors are provided in parentheses. Restriction sites are underlined. SsfA and SsfB genes were synthesized in a single vector to yield a dicistronic plasmid.

| Name | Sequence |
|---|---|
| SsfA-SsfB<br>co-expression<br>(pET15b) | CATATGTTCGTGCATAGTGATCGTTTTCCGGTTAGTAGCCCGCTGCCGAGCGGCGTTCC<br>GACCCCTGTTCCGTGGGGCGTTAGTCGCATGGCACCGTATCCGACCACCGCACCGGAT<br>TATGCACGTGTTGAACTGGACCCTGATACCCAGACCGCCGTGTTTTTGATAAAGATGG<br>TCAGGTGATGGAAATGGGCAAACATAATACCAGTACCGGTACCAGTCCGGCAACCGGT<br>ACCTCACCGGATGGTAATGGTACCACCCAGGATAGCGATACCGGCAGTGATAGTGATC<br>AGTAAtctagaaataattttgtttaactttaagaaggagatataccATGGGCAGCAGCCATCATCATCATCATC<br>ACAGCAGCGGCCACATGAGCGAACCGCGTCCGGTGCTGGTTGTGACCAGCCTGCATGA<br>TCCGACCGCCGATGTGGTGATTCGTGAACTGCATGATCGTGGCGTTCCGGTTGTTCGTC<br>TGGATAGTGGTGATTTTCCGGCAAGTCTGAGTGTTGAAGCAGAAATTACCAGCAGCGGT<br>CTGCGTGGTCGCCTGCGTACCCTGACCCGTACCGCAGATTTAGCAGGCGTGCGCAGCC<br>TGTATTATCGTCGTCCGACCGGCTTTGCATTTCCGCATCTGGATGAACAGGATGCACGT<br>TATGCAGTTACCCAGGCCCGTTATGGCCTGGGCGGTGTGCTGGCCAGTCTGCCTGGCT<br>GCCTGTATGTGAATCATCCGCATCGCGTGGGTGATGCAGAATTTAAACCGAGTGGCCTG<br>GCAGTGGCAGTTAGTGCAGGCTTTCTGGTTCCGCCGACCCTGATTACCAGCTCACCGG<br>ATGCAGCCCGTGCCTTTATTAAACAGCATGGCCCGGTGATTTATAAACCGCTGTATAATC<br>CGGTGTATCGCATTGATGGCGTTAGCAGCGTTGTTAAAATTGCCGAAGTGGCAGTTGAA<br>GATATTGATGATAGTGTTGCAGGTACCGCCCATCTGTTTCAGCAGCGTGTGGAAAAAAT<br>TAGTGATGTTCGCGTTACCGTGATTGGCGATCAGGTGTTTGCCGTGCGTATTGATAGCG<br>ATCTGCTGGATTGGCGCACCGATTATAGTCGCCTGCGTTATAGCGTTGATACCCCGCCG<br>CCGGGCGTTACCGAAGCCTTACATGCCTATCTGAGTCATTTTGGTCTGGTGTTTGGTGC<br>ATTTGATTTTGCCGTTGATCGCGCCGGTCGCTGGTGGTTTCTGGAATGTAATCCGAGTG<br>GCCAGTGGTATTGGCTGGAAGATGAAACCGGTCTGCCGATGCTGGCCGCAATGGCCGA<br>TCTGCTGGAACGTAAAAGCCTGTAACTCGAG |
| SsfM<br>(pCDFDuet-1) | GGATCCGATGACCGTGATGGATAGCGCCGCCCTGCGTCGCGAACTGGCAGGTCGTCT<br>GGCAGCAGGCGGTAGCCTGCGCACCGAACCGTGGCGTAAAGCAGTGGAAGCCGTGCC<br>GCGCCATGAATTTCTGCGTGGTGGCTTTTTTGTGCGTAGTGCACCGGATGCCTGGCGTC<br>CGGTGCTGGCAGATGCAGATGGCTGGCTGGAACGCTGTTATGCCGATGAAAGTCTGGT<br>TACCCAGATTGCCGGCACCATTGTTCCGGGTGATGTGCGTGGTGAAATTCTGCGTGAAC<br>CGACCAGCAGTAGCACCCTGCCGAGCCTGGTGGTGCGCATGATGGAAGATTTACAGGT<br>TGAAGAAGGTGATCGTGTGCTGGAAATTGGCACCGGTACCGGTTATAGTACCGGCCTG<br>CTGTGCCATCGTCTGGGTGATGATCTGGTTACCAGTGTGGAAGTTGATGCAGAAGTGAG<br>TCGCCGTGCAGGTACCGCCCTGGGCGCATGTGGCTATTTTCCGGAACTGGTGGTGGGC<br>GATGGTCTGGCAGGTCATAAAGATGGTGCACCGTATGATCGTCTGATTGCCACCTGTGG<br>TGTTCGTCAGCTGCCGTATACCTGGATTGAACAGACCAAACCGGGCGGCGTTATTCTGG<br>CAACCGTGTGCGGCTGGATGCATGCAAGCGAACTGGCACGTCTGACCGTGGGTGATGA<br>TGGCGAAGCCCGCGGCCGCCTGCTGGGTGGTCAAGTTAGCTTTATGCCGGCCCGTCC<br>GCAGATGCCGCCGCCTTTAGGTATGCTGCCGGATCTGGAAAGCGGTCAGGAACGTGTT<br>GCCCGTTTTGGTCCGGAAGTGCTGGATGCCTGGGATGCCCGTTTTGTGGCCCAGCTGG<br>CCGCCCCTCATGCCCAACGTCTGAGCCTGACCCTGGCAGATAAAACCCAGCATGTGCT<br>GATTGATGTTGAAGCCGGCGCCTGGGCAGCACTGGTTCTGGATGGTGAACATCGCACC<br>GTGCGCCAGGGTGGCCCGGTTCAACTGTGGGATCAGATTGAAGAACATCTGCTGCGTT<br>GGCAGGCCGATGGTAGTCCGGCCCTGACCCGCTTTGAAATTCTGGTGACCAGTGAAGG<br>CCAGCATTTTACCTGGCCGAAAAGCTAACTCGAG |
| FcaA<br>(pET22b) | CATATGGGTAGTAGCCATCATCATCATCACCATAGTAGTGGCCACATGGGCAAACATGC<br>CAAACCGGTTGCATGTCCGACCTGTAATGGTAGCGGTAAAATTACCGTTACCAGCGATG<br>GTAAAAATGAAACCGTTAGCTGTGGCGTGTGCAAAGGTAGCGGTAAAGCATAACTCGAG |

| | |
|---|---|
| FcaM<br>(pCDFDuet-1) | GGATCCGATGAGCGTTCGCCGCGTGAATCGCGAAGATTTTATTCCGGATGAAATTTGGG<br>TTCGCGATGAAGATGGTTTCTTTCTGGTGCCGCTGCGTCGCCAGGATGATCCGCAGCG<br>TTGGAGTGAACTGTGCCGTGGCGATGATGGTATTACCACCCAGGTTGATGATGGTACCG<br>GCCGTTATGATGGTCGTGGCGTGATTCCGACCAGTAGCAGTAGTGCCCCGTGGGTTAT<br>GGCCCGCATGCTGGATCTGCTGGATGTGCGCGATGGCATGAATGTGCTGGAAATTGGC<br>ACCGGTACCGGTTATAATGCAGCCCTGCTGGCCGAACGCACCCCGACAGGTCAGGTTA<br>CCACCATTGAAATTGATCCGGGTATTGCCGGCCATGCACGCGCCGTGCTGGCAAGAAT<br>TGGCCGTCCGGTGACCGTGGTTGTGGGCGATGGTGCCGCCGGTTTTCCGGATCGTGC<br>ACCGTATGATCGCATTATTGCCACCGCCAGCGTTGTGACCGTTCCGTATCCGTGGATTA<br>CCCAGACCCGCCCGGGCGGTCGTATTGTGCTGCCGTTTACCAGTGAATTTGGTGGCGC<br>ACTGCTGAGTCTGACCGTGGCAGATGGCACCGCCAGTGGTCATTTTCATGATGATGCAG<br>GCTTTATGCGCCTGCGTGGCCAGCGCGCAGATCGTCCTGTGTGGTGGCTGGGCGAAG<br>ATGATGCCGATGTGCGTACCACCTGTCGTTATCTGGATGAACCGTTTGCCGATGCAGCC<br>GCCGGCTTTGCCGTGGGTCTGTGGCTGCCGGGCTGCACAACCGGCCAGATTGAAGAA<br>GGCGGCCCGGCACGCACCCTGCTGTTAAGCCATGCCGCAAGTCATAGTTGGGCAAGTC<br>TGACCGCCGGCAGTGATGGCCAGAGTGATGGCCATGAAGTTACCCAGTATGGTCCGCG<br>TCGCCTGTGGGATGAACTGGAAATGGCATATGATTGGTGGGTGAATGCCGGCCGTCCG<br>AGCTGTGCCCGTTTTGGTCTGACCGTTACCCCGGATGGCCAGACCAGTTGGCTGGATT<br>ATCCGGAACGCGTGATTCCGGCCAGCTAACTCGAG |
| SpaA<br>(pET22b) | CATATGGGTAGTAGTCATCATCATCATCACCATAGCAGTGGTCACATGGCAGGCAAACA<br>TGAAAAACCGGAACCGGACCCTGGCCAGGGCGCACCTCCTCCTGGTAATAGCGATGGC<br>CAGGTTCCGCCGCCGCCGCCTGGTAATGGCAAACGTAAAAAATAACTCGAG |
| SpaM<br>(pCDFDuet-1) | GGATCCGATGGAATACGATGAACAGCGCGGTCGCCTGGCAGAAGTGATGGATGAACGC<br>GGTGCCTGGCCGGCACGCAGCCCTTGGATTCGCCAGGCCGTTGATGCACTGCCGCGC<br>CATCATTTTGCACCGGATCGCCTGTGGCGCTGGAATGGTCATGCCTATATTCCGGTTGA<br>TCGCGATGCAGATCCGGAACAGTGGGCCGCAGAAGTTTATGCAAGTCCGGATACCGCC<br>GCAGTTACCCAGGTTACCGATGGTGTTCCGAGCAGCAGCCTGAGTGCACAGGGCGTTG<br>TGGTTGATATGCTGGATAGCCTGCTGGTTGAACCGGGTGATCGCGTTCTGGAACTGGG<br>TGCAGGTACCGGCTGGAATGCAGCCCTGCTGGCAGAACGTGCCGGCCCTGGTCGCGT<br>TGTTAGCGTTGAAGTGGATGCCGAACTGGCAGCCGCCGCCGCTGGTAGACTGAAAGCA<br>GCCGGTGCCGATGTGGCAGTGGAAATTGGTGATGGTGCCGCAGGTCGTCCGAGTGGT<br>GCCCCTTATGATCGCCTGATTAGTACCTATGCCGTTAGCACCGTGCCGTGGGCATGGGT<br>GGAACAGACCCGCCCGGGTGGTCGCATTGTTACCCCGTGGGGTCGTATGGGTCATGTT<br>GCCCTGACCGTGGCAGATGATGGTCGCAGTGCCACCGGTTGGGTGCAGGGCCTGGCC<br>ATGTTTATGCCGGCCCGTGGTACCCGCAGCGCCTTAGGTTGGCGTCAGATTCATGGTG<br>ATGGCCCGCCGGATGATGAACGTCCGTTTGGTCGTGATCTGCGCCCGCTGCGCGAAGA<br>TGCCCATCTGCTGTTTGCCCTGCGCGTGCAGCTGCCGGATGTGCGTGTGACCACCGGC<br>GTTGATGATGATGGCGTGAATGTTTGGCTGCATGATGGTGTTGCAAGCTGGGCCACCCT<br>GACCACCCTGCCGGATGGTCAGGCCACCGCAAGCCAGGGCGGCCCTAGACGTCTGGC<br>CGATGAACTGGAACCGGCATGGGATTGGTGGGTGAGCGAAGGCGAACCGACCCTGTAT<br>GATTTTGGCATGACCGTGGAACCGGATCGTCAGTATGTTTGGTGTCGCGATGCAGCAAC<br>CGGCCCGCGCTGGCCTGTGGAAGCTATGCCGTAAAGATCT |

**Table S2. Plasmids used in this study.**

| Plasmid | Vector | Protein | Reference |
|---|---|---|---|
| pHB388* | pET22b | His6_MBP_[TEV]_PsnA2 leader-(LVPR)-PsnA2 core | [29] |
| pHB665 | pET15b | $His_6$_[Tm]_SsfA, $His_6$_SsfB | This study |
| pHB684 | pET15b | $His_6$_[Tm]_SsfA | This study |
| pHB666 | pCDFDuet-1 | $His_6$_[Tm]_SsfM | This study |
| pHB763 | pET22b | $His_6$_FcaA | This study |
| pHB772 | pET22b | $His_6$_MBP_[TEV]_FcaA | This study |
| pHB764 | pCDFDuet-1 | $His_6$_FcaM | This study |
| pHB717 | pET22b | $His_6$_SpaA | This study |
| pHB727 | pET22b | $His_6$_MBP_[TEV]_SpaA | This study |
| pHB718 | pCDFDuet-1 | $His_6$_SpaM | This study |

[Tm] = Thrombin cleavage site
[TEV] = TEV protease cleavage site
* This plasmid was used as template for the cloning of MBP-fused precursors

**Table S3. Oligonucleotides used in this study.**

| Primer | Sequence | Description |
|---|---|---|
| T7P | TAATACGACTCACTATAGGG | For sequencing |
| T7T | GCTAGTTATTGCTCAGCGG | For sequencing |
| F_Stop | CTCGAGGATCCGGCTGCTAACAAA | Deletion of SsfB gene in pHB665 |
| R_SsfA | TTACTGATCACTATCACTGCCGGTATCGCT | Deletion of SsfB gene in pHB665 |
| F_Amp | GACACCACGATGCCTGCAGCA | Amplification of MBP and [TEV] containing DNA fragment in pHB388 |
| R_TEV | GCCCTGGAAATACAGGTTTTCTCCGGAGCT | Amplification of MBP and [TEV] containing DNA fragment in pHB388 |
| R_Amp | GCGCAACGTTGTTGCCATTGC | Amplification of FcaA or SpaA gene in the corresponding plasmid |
| F_FcaA | ATGGGCAAACATGCCAAACCGGTT | Amplification of FcaA gene in pHB763 |
| F_SpaA | ATGGCAGGCAAACATGAAAAACCGGAACCG | Amplification of SpaA gene in pHB717 |

[TEV] = TEV cleavage site (ENLYFQ/G; gaaaacctgtatttccagggc)
